# Flexible machine learning prediction of antigen presentation for rare and common HLA-I alleles

**DOI:** 10.1101/2020.04.25.061069

**Authors:** Barbara Bravi, Jérôme Tubiana, Simona Cocco, Rémi Monasson, Thierry Mora, Aleksandra M. Walczak

## Abstract

The recent increase of immunopeptidomic data, obtained by mass spectrometry or binding assays, opens unprecedented possibilities for investigating endogenous antigen presentation by the highly polymorphic human leukocyte antigen class I (HLA-I) protein. We introduce a flexible and easily interpretable peptide presentation prediction method, RBM-MHC. We validate its performance as a predictor of cancer neoantigens and viral epitopes and we use it to reconstruct peptide motifs presented on specific HLA-I molecules. By benchmarking RBM-MHC performance on a wide range of HLA-I alleles, we show its importance to improve prediction accuracy for rarer alleles.

## Introduction

Recognition of malignant and infected cells by the adaptive immune system requires binding of cytotoxic T-cell receptors to antigens, short peptides (typically 8-11-mers) presented by the Major Histocompatibilty Complex (MHC) class I coded by HLA-I alleles (Fig. 1A). Tumour-specific neoantigens, *i.e*. antigens carrying cancer-specific mutations, are currently sought-after targets for improving cancer immunotherapy [1, 2]. Computational predictions can help select potential neoantigens and accelerate immunogenicity testing. To be useful, these predictions must be specific to each HLA type.

**Figure 1:**
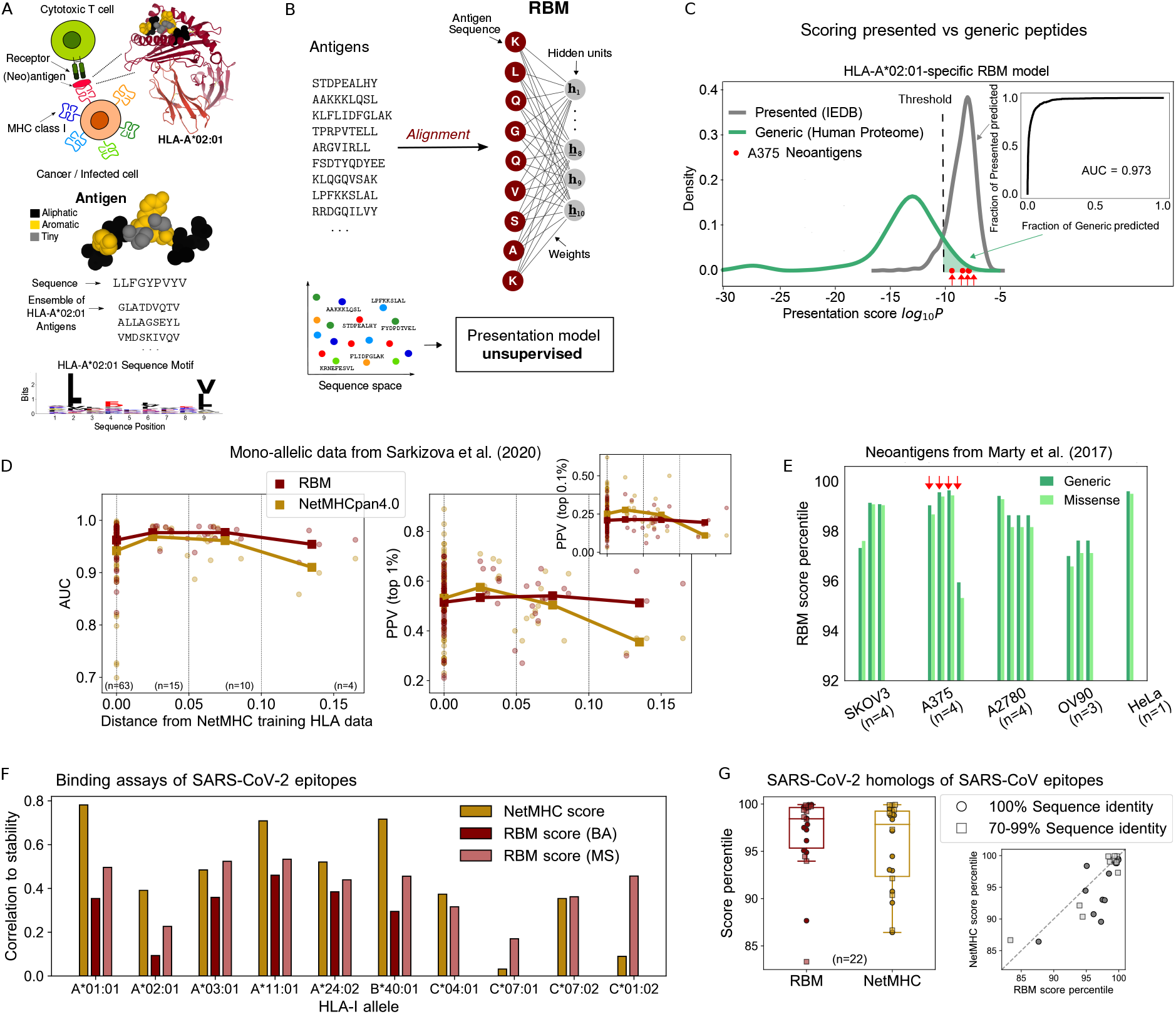
RBM approach to HLA-I antigen presentation. (A) Antigens binding to a given HLA match specific “sequence motifs” represented by logos (example of Tax-HLA-A*02:01 complex structure, PDB-ID:1BD2, Mol* image [33]). (B) RBM model structure. (C) RBM scores for presented *vs* generic peptides on HLA-A*02:01 and predictive performance assessed by ROC. (D) RBM and NetMHC performance at recovering MS-detected peptides from [6] for 92 HLA-I alleles as measured by AUC and PPV metrics. PPVs and AUCs are plotted as a function of the distance between the corresponding HLA and the closest one in NetMHCpan4.0 training dataset. Bold lines highlight the trend of mean AUC and PPV values (plotted by squares) over subsets of alleles grouped by distance as indicated by vertical dashed lines (63 alleles with distance = 0, 15 alleles with 0 < distance < 0.05, 10 alleles with 0.05 < distance < 0.1, 4 alleles with distance > 0.1). (E) Score percentiles on the same allele as (C) of the neoantigens from 5 cancer cell lines validated in [23] - 4 neoantigens validated for cell line SKOV3, 4 for A375, 4 for A2780, 3 for OV90 and 1 for HeLa. (F) Correlation of experimental stability of SARS-CoV-2 epitopes (94 for each HLA allele [24]) with scores from NetMHC4.0/NetMHCpan4.0 and scores from RBM trained on Binding Affinity (BA) - not available for HLA-C alleles under consideration - and Mass Spectroscopy (MS) data (with *ad hoc* sequence re-weighting), see also STAR Methods and Supplementary Fig. 6B. (G) RBM and NetMHC4.0 score percentiles (relative to all SARS-CoV-2 9-mers) of *n* = 22 homologs of dominant SARS-CoV epitopes identified in [25]. Boxplots indicate median, upper and lower quartiles.

State-of-the-art methods [3–6], such as NetMHC [7–9], are based on artificial neural networks trained in a *supervised* way to predict peptide presentation from known peptide-HLA association. They are trained on large datasets at every method release and they provide peptide-scoring schemes that perform best on frequent alleles and well characterized alleles at the time of release. Their accuracy is degraded for rare or little studied HLA-I alleles which are poorly represented in databases. In that case, another approach is to train *unsupervised* models of presentation from custom elution experiments with little or no information about peptide-HLA association. For instance, MixMHCp [10–12] can reconstruct, from unannotated peptide sequences, a mixture of generative models—one for each expressed HLA type. However, it makes simplifying assumptions about binding specificity, and is not designed to leverage available (albeit limited) annotation information from the Immune Epitope Database [13] (IEDB) to improve accuracy.

We present an alternative method, called RBM-MHC, for predicting peptides presented by specific class I MHCs. RBM-MHC is a scoring and classification scheme that can be easily trained “on the fly” on custom datasets, such as patient- or experiment-specific samples, and more generally on newly available data. As such, RBM-MHC enables improved predictions for rare alleles at a fast pace, without waiting for new releases of general software like NetMHC [7–9], with which it is not intended to compete on an equal footing.

The method consists of two parts. The first component (Fig. 1) relies on a Restricted Boltzmann Machine (RBM), an unsupervised machine learning scheme that learns probability distributions of sequences given as input [14–16]. The RBM estimates presentation scores for each peptide, and can generate candidate presentable peptides. The RBM also provides a lower dimensional representation of peptides with a clear interpretation in terms of associated HLA type. The second component of the method (Fig. 2) exploits this efficient representation to classify sequences by HLA restriction in a supervised way, using only a small number of annotations.

**Figure 2:**
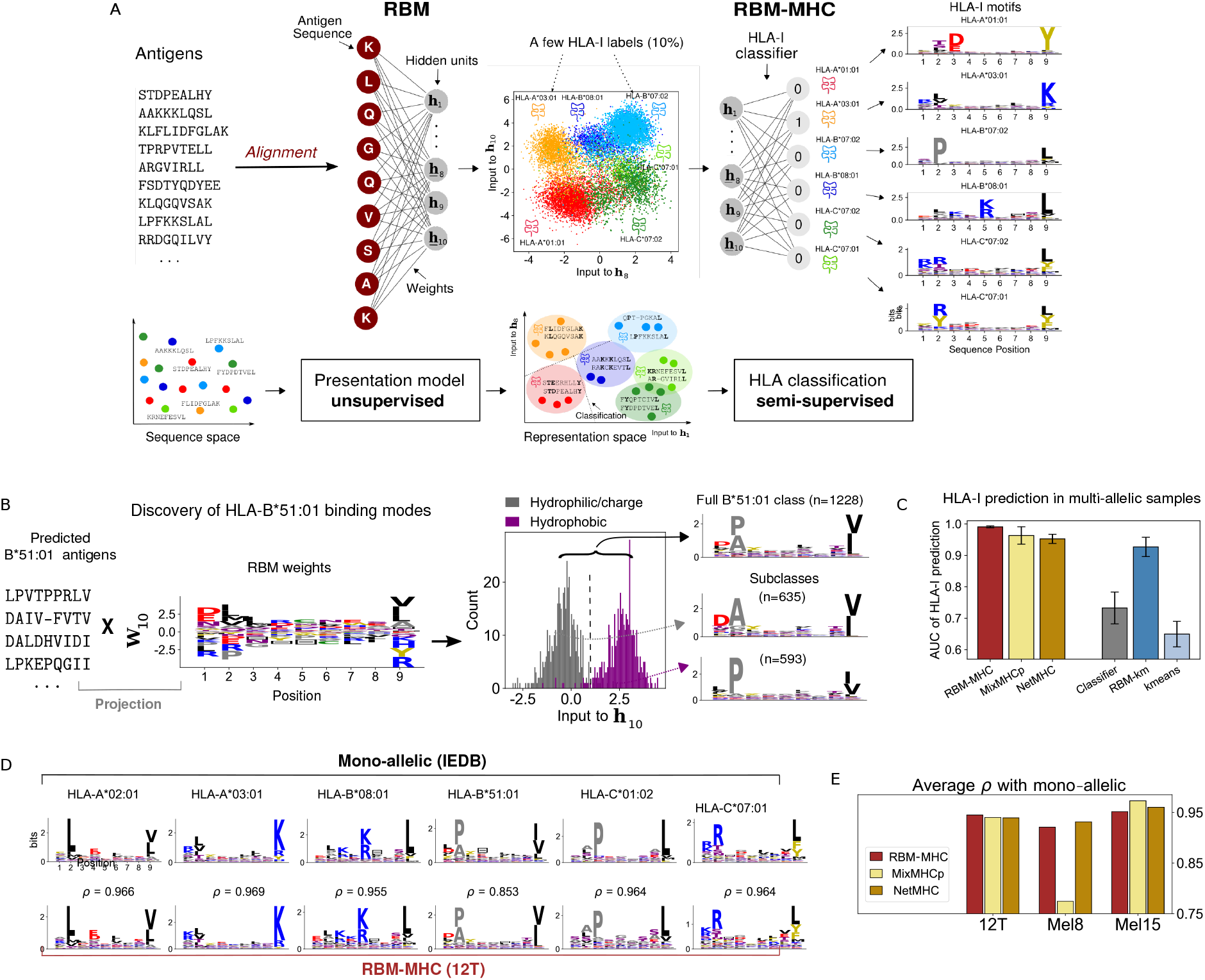
RBM-MHC approach to HLA-I classification. (A) RBM-MHC workflow. RBM projects sequences onto a representation in terms of “hidden units” (we selected hidden unit 8 and 10 for illustration) through the set of learnt weights. In this representation space, each cluster groups together antigens with the same HLA-binding specificity (given by the color code). Linear classification, guided by the knowledge on the HLA-I restriction of a few antigens in each cluster (“labels”), is performed through the HLA-I classifier to predict the HLA-I type of all antigens. Data: IEDB-derived dataset for Haplotype 1, Supplementary Table 2. (B) RBM distinguishes 2 subclasses in HLA-B*51:01-binding antigens. The HLA-B*51:01-peptide bond can be established [12]: *(i)* via the interaction of the HLA-B*51:01 residue 62 with peptide position 2, requiring a hydrophilic residue there, typically Alanine (A), and a polar or negatively charged residue, typically Aspartic Acid (D), at position 1; (ii) with R62 sidechain facing the solvent, requiring co-occurrence at positions 1-2 of hydrophobic residues, typically Proline (P), at position 2). Inspection of the inputs to the 10th hidden unit (**h**_10_), found by projecting peptides predicted by RBM-MHC as HLA-B*51:01-specific onto the corresponding weights (STAR Methods, Eq. 4), reveals a bimodal distribution, enabling the discrimination of the “hydrophilic/charged” pattern *(i)* from the “hydrophobic” pattern *(ii)*, as recapitulated by the sequence logos. (C) Performance (AUC) of 6 methods for HLA-I prediction on 9-mers in 10 synthetic-individual samples, each carrying 6 HLA-I covering A,B,C alleles (see Supplementary Table 2). Bars are standard deviations over the 10 datasets. (D) Sequence logos of clusters found with RBM-MHC trained on melanoma-associated sample 12T [26, 27] and IEDB mono-allelic data; *p* gives the Pearson correlation between respective amino acid frequencies. (E) Performances (average Pearson correlation p over clusters) for 3 samples from [26, 27, 31]. In C,E the MixMHCp and NetMHC versions used are respectively MixMHCp2.1 and NetMHCpan4.1.

## Results

### Restricted Boltzmann Machine for peptide presentation

The first building block of the method is an RBM, a probabilistic graphical model defined by a simple structure with one hidden layer and weights connecting the input—the peptide sequence—to the hidden layer (Fig. 1B). RBM parameters, *i.e*. the weights and the biases acting on both input and hidden units, are learned from a training set of presented peptides collected from public repositories or custom samples (STAR Methods, Supplementary Figs. 1A, 2A and 3). These input peptides may come from mass spectrometry (MS) experiments or binding assays. They may come from several HLA alleles, or from a single one. For example, in datasets from single-individual samples, the number of HLA is at most 6, as each individual inherits 2 alleles of each locus HLA-A, HLA-B and HLA-C. Another example is given by mono-allelic samples [4, 6], where peptides are presented by a single HLA protein.

After training, the RBM returns for each sequence a probability, which is interpreted as a global score of antigen presentation by the HLA proteins involved in the training dataset. Since learning the RBM requires fixed-length input sequences, we first reduce peptides of variable length to a reference length by an alignment procedure based on a Hidden Markov Model profile built from aligning subsets of same-length sequences using standard routines (STAR Methods and Supplementary Fig. 1E). We set the reference length to 9 residues, the most common length of peptides presented by class I MHCs [6, 7, 17, 18].

### RBM performance on mono-allelic data

As a first validity check of RBM-based presentation scores, we built a RBM model to predict presentation by a single common allele, HLA-A*02:01, from MS data annotated with this HLA restriction in IEDB [13]. We tested the model score’s ability to discriminate presented peptides from “generic” peptides (randomly drawn from the human proteome). We calculated a Receiver Operating Characteristic (ROC) curve from RBM presentation scores assigned to a test set of presented versus generic peptides (Fig. 1C). The Area Under the Curve of the receiving operating characteristic, AUC = 0.973, by far above the random-expectation value (AUC = 0.5) and close to the maximal value (AUC = 1), proves the RBM predictive power at recovering presented antigens.

We next benchmarked RBM’s predictive power on recent MS datasets from mono-allelic cell lines [6]. In addition to the AUC, which considers with equal weight presented and generic peptides, we computed the Positive Predictive Value (PPV) [4, 6], which measures the model’s ability to correctly recognize presented peptides among a 99- or 999-fold excess of generic peptides (equivalent to assuming that only 1% [19–21] or 0.1% [4, 6] of all peptides bind to a given HLA, see STAR Methods). Supplementary Fig. 5B compares the AUC and PPV between RBM and NetMHCpan4.0 [8] for all the 92 different HLA alleles encompassing A, B, C loci from [6] (31 HLA-A, 40 HLA-B, 21 HLA-C). NetMHCpan4.0 is the penultimate version of NetMHCpan, which was trained before the publication of the mono-allelic data [6] and is therefore trained on an independent dataset. In contrast RBM was trained on data presently available in IEDB for all the 92 alleles. To ensure independence from the training set, testing peptides were manually excluded from training data (STAR Methods). Both methods performed comparably (AUC = 0.97 for RBM *vs* 0.95 for NetMHC, 1% PPV = 0.52 for RBM *vs* 0.53 for NetMHC).

When there is limited or no data for a given HLA allele, NetMHCpan extrapolates from the most similar HLA allele for which data are available. Fig. 1D shows that NetMHCpan performance degrades with the distance between the queried HLA and that nearest neighbor in the training dataset [22]. By contrast, the RBM performs equally well on all HLAs, which is expected since in that case the distance to the training dataset is always zero. This highlights the importance of being able to flexibly and rapidly train models on new data, especially for HLA alleles that are poorly covered in previously available datasets. The four alleles at largest distance from NetMHCpan4.0 training data, for which RBM outperforms NetMHC, are all HLA-C (HLA-C*01:02, HLA-C*02:02, HLA-C*07:04, HLA-C*17:01, see Supplementary Fig. 5B), a locus usually under-represented in existing databases.

### RBM can predict cancer neoantigens

An important application of antigen presentability is the identification of neoantigens arising from cancer mutations, which are key to evaluating their potential for immunotherapies. To assess RBM’s ability to predict neoantigens, we looked at missense mutations in 5 ovarian and melanoma cancer cell lines from Ref. [23]. We attributed presentation scores to all 8-11-mer peptides harboring these mutations, using the RBM previously trained on HLA*A:02:01 (known to be expressed in the 5 cell lines), and also computed, for comparison, the corresponding score by NetMHCpan4.1 [9], the latest version of NetMHCpan. 16 neoantigens were experimentally validated by MS to associate with HLA*A:02:01 [23]. Of those, 15 were ranked by RBM in the top 4.1% among generic peptides (mean score percentile 1.5% versus 1.6% for NetMHC) and 4.7% among mutated peptides (mean 1.8% versus 0.9% for NetMHC) for the corresponding cell line. This demonstrates that the RBM reaches state-of-the-art performance at predicting presented neoantigens.

### SARS-CoV-2 epitope discovery

Predicting which antigens are presented by virus-infected cells is key to the rational design of vaccines targeting these antigens. We tested RBM’s ability to perform this task on the example of SARS-CoV-2, using recent *in vitro* measurements of binding stability [24] of SARS-CoV-2 candidate epitopes (selected using NetMHC4.0 [7] and NetMHCpan4.0 [8], see STAR Methods).

We first trained an allele-specific RBM for each HLA-I involved in the experiment. Since the experiment measures binding, we chose our training sets as binding assay (BA) datasets, as well as MS datasets that were reweighted to make them comparable to BA (correcting for amino-acid frequency biases, see STAR Methods). We then attributed an RBM score to each peptide, and compared it to its experimental binding stability (Supplementary Fig. 6A-B). In Fig. 1F we report the correlation between the RBM score and measurement for each allele. The performance of the NetMHC version used to select the epitopes (NetMHC4.0/NetMHCpan4.0) is shown for comparison. As before, RBM outperforms NetMHC4.0/NetMHCpan4.0 for rarer HLA-C alleles (HLA-C*07:01 and HLA-C*01:02). It is noteworthy that correlations scores for rare alleles are improved using the very recent NetMHCpan4.1 and get comparable to RBM results (see Supplementary Fig. 6C). Supported by the good performance of RBM on rare alleles, we suggest new SARS-CoV-2 epitopes for HLA-C alleles that were not among the top-scoring ones by NetMHC4.0/NetMHCpan4.0. Our predictions, given in Supplementary Table 1, have been favorably cross-checked with NetMHCpan4.1 scores.

RBM also assigns high presentation scores to SARS-CoV-2 epitopes that are homologous to experimentally validated SARS-CoV cytotoxic T-cell epitopes [25], on par with NetMHC4.0 (Fig. 2F) and NetMHCpan4.0/NetMHCpan4.1 (Supplementary Fig. 6E).

### RBM offers useful low-dimension representations

In addition to providing a presentation score, RBM hidden units allow for mapping peptides onto a lower dimensional “representation space” given by the inputs to the RBM hidden units [16] (see STAR Methods). The full potential of this representation is best illustrated on an RBM trained on multi-allelic data, where peptides may be bound to up to 6 different HLA-I proteins. Fig. 2A shows an example of such a representation projected in 2 dimensions, on a synthetic dataset obtained by pooling peptides from 6 HLA restrictions in IEDB. The low-dimensional projection organizes peptides into 6 well-defined clusters, reflecting the HLA-binding specificities present in the sample. Each HLA recognizes specific “sequence motifs” (patterns of preferred residues at each position) in the presented peptides. The data-derived RBM parameters underlying that representation play a key role in capturing what amino acids contribute to define HLA-binding motifs, as illustrated on a bi-allelic case in Supplementary Fig. 2B.

The representation space can also be useful to reveal new features even in the mono-allelic case. A compelling example is provided by HLA-B*51:01-restricted peptides derived from a clinical sample [26, 27] (see below for details about how restriction was predicted). Projection of HLA-B*51:01-specific sequences onto a single RBM feature reveals a double-peaked histogram, corresponding to two structurally alternative binding modes, which were validated in [12] (Fig. 2B).

### HLA classification with RBM-MHC

This low-dimensional representation of peptides suggests an efficient way to classify them by HLA-I specificity, using the annotation of a small number of sequences by their HLA preference. In practice these annotated sequences may come from other experiments or public databases as IEDB [13]. Using these annotations, we train a linear classifier to predict the HLA restriction of individual peptides from their RBM representation (see STAR Methods, Fig. 2A, Supplementary Figs. 2A, 3C-E). We refer to this architecture, which combines the RBM (trained on unannotated peptides) *and* the HLA-I classifier (trained on a few annotated ones), as RBM-MHC.

To test performance in a case where the ground truth is known, we trained RBMs on 10 synthetic “single-individual” samples pooling together IEDB 9-mer antigens presented by 6 HLA-I proteins per individual, covering 43 different alleles in total (12 HLA-A, 17 HLA-B, 14 HLA-C, see STAR Methods and Supplementary Table 2). We randomly selected 10% of peptides and labeled them with their associated HLA, and used them to train the RBM-MHC classifier. The performance of RBM-MHC at predicting HLA association, as measured by AUC = 0.991, is excellent (Fig. 2C and STAR Methods). The RBM representation is crucial for achieving high prediction performance. Training a linear HLA-I classifier directly on the annotated sequences (rather than their RBM representation) yields a much poorer performance (AUC=0.733). On the other hand, a completely unsupervised clustering algorithm (K-means [28]) applied to unlabeled data in RBM representation space (“RBM-km”) performed well (AUC=0.927), while K-means applied directly to sequences did not (AUC=0.650, Fig. 2C and STAR Methods).

The structure of the RBM representation space further allows for setting up a well interpretable protocol to generate new, artificial peptides from the model with controled HLA-binding specificity (STAR Methods, Supplementary Fig. 2C-E).

We next extended our HLA type predictions to peptides of variable length (8-11 residues). Multi-allelic datasets pose the additional challenge that sequences must be aligned together across different HLA motifs. To optimize this multiple sequence alignment, we resorted to an iterative procedure, where the sequence alignment is refined using the the HLA assignments from a first iteration of RBM-MHC, after which a second RBM-MHC is trained to obtain the final model. This procedure allows the steps of HLA classification and alignment to inform each other (STAR Methods and Supplementary Fig. 1E). We benchmarked our alignment routine against other well established alignment strategies (based on the MAFFT software [29]) and we verified that our routine ensures higher HLA classification performance, even without the refinement step (STAR Methods and Supplementary Fig. 1F-L).

### RBM-MHC compares favorably to existing methods

RBM-MHC outperforms MixMHCp2.1 [10–12] in terms of both overall accuracy and stability across the 10 synthetic datasets (Fig. 2C, Supplementary Fig. 7). This is in part thanks to the 10% labeled data it exploits. The RBM also captures global sequence correlations that MixMHCp’s independent-site models miss, which are important for correctly classifying antigens across alleles with similar binding motifs. The major drop in MixMHCp2.1 performance occurs precisely in datasets that mix same-supertype alleles [30] (HLA-B*27:05 and HLA-B*39:01 belonging to supertype B27, HLA-B*40:01 and HLA-B*44:03 belonging to B44, see Supplementary Table 2). RBM-MHC also performs better in terms of AUC than NetMHCpan4.1 [9], which can be applied here to predict binding to each of the 6 alleles in our datasets, and stays comparable to NetMHCpan4.1 when several other performance indicators are considered (Supplementary Fig. 7). The gain in AUC of RBM-MHC over MixMHCp and NetMHC is slightly more pronounced for 9-mers-only samples than for 8-11-mers (Supplementary Fig. 7A), suggesting room for further improvement in the alignment.

### Application to single-patient cancer immunopeptidome

To assess the relevance of our approach in a clinical setting, we considered single-patient, melanoma-associated immunopeptidomic datasets [26, 27, 31], complemented with patient HLA typing and Whole Exome Sequencing (WES) of tumorous cells, where in total 11 neoantigens were identified. We tested the RBM-MHC approach for motif reconstruction. Since in this case true peptide-HLA associations are unknown, model performance is evaluated through the correlation between the predicted motifs and motifs reconstructed from IEDB mono-allelic data (Fig. 2D-E, Supplementary Fig. 8). RBM-MHC, MixMHCp2.1 and NetMHCpan4.1 perform comparably, with the only exception of sample Mel8, where MixMHCp2.1 merges antigens specific to the 2 HLA-C into the same cluster, causing a drop in the average *ρ* (Fig. 2E, Supplementary Fig. 8B). In addition, RBM-MHC and its unsupervised version RBM-km trained on the patient dataset (see STAR Methods) systematically predicted the correct HLA association for all neoantigens. They also assigned a top 1% score among all WES mutants to 8 out of 11 identified neoantigens (Supplementary Table 3).

## Discussion

In this work, we presented RBM-MHC, a novel predictor of antigen sequences presented on specific HLA-I types. It can be broadly used for 4 tasks. First, the RBM module of the method can be used to score the presentability of (neo-)antigens, regardless of HLA information (Fig. 1). Second, the low-dimensional representation of peptides provided by the RBM allows one to visualize different HLA binding motifs (as in Fig. 2A), or different binding modes within the same HLA (as in Fig. 2B), providing a useful tool for data exploration and feature discovery. Third, RBM-MHC may be used to classify peptides by HLA restriction using only a limited number of annotations, matching state-of-the-art performance (Fig. 2C). Finally, the method can generate putative antigens with or without a specific HLA restriction (Supplementary Fig. 2C-E). These generated peptides could be experimentally tested in future, providing additional evidence for the validity of the model [32].

RBM-MHC’s use will also depend on the choice of training dataset. Single-allele datasets allow for training allele-specific models to score antigen presentation by a specific HLA allele. In this way, we have proposed new putative SARS-CoV-2 epitopes for HLA-C alleles that could be tested experimentally (Supplementary Table 1). Multi-allelic datasets (from *e.g*. peptidome mass spectrometry of clinical samples), allow for training models to score antigen presentation in a given donor (with all its HLA-I alleles), and to predict the HLA-binding specificity of given peptides. We benchmarked our approach on several examples of datasets. In single-allele datasets, we found that RBM-MHC can perform similarly to established methods, like NetMHCpan, for frequent, well-characterized alleles (Figs. 1D-G, Supplementary Figs. 5 and 6). This finding is remarkable, since NetMHC training is fully supervised, incorporates more information such as binding affinities, and uses positive (binders) as well as negative (non-binders) examples. The number of binders per allele in the version NetMHCpan4.0 ranges between ~ 50 and ~ 9500 [8], while RBM-MHC is trained on datasets of presented peptides only, making our approach less data-demanding. The amount of 8-11mer peptides per allele considered in this work from IEDB-derived MS records varies from ~ 500 to ~ 10^4^, with an average (2350) comparable to NetMHCpan4.0 positive examples. RBM-MHC is a more flexible machine-learning scheme that can be easily trained on newly-produced datasets, tracking the fast growing number of available datasets to improve its predictive power, especially for previously under-represented HLA-I alleles (Figs. 1D,F, Supplementary Fig. 5), such as HLA-C alleles.

The latest version of NetMHCpan, NetMHCpan4.1 [9], was published very recently, 3 years after the previous release NetMHCpan4.0 [8]. This new version was trained on a total of ~ 850,000 ligands collected from public (mainly IEDB) and in-house resources [9]—a 10-fold increase with respect to NetMHCpan4.0. All our comparisons were done with this latest version, except on the mono-allelic data from Ref. [6] (Fig. 1D) and on the SARS-CoV-2 epitope data from [24] (Fig. 1F), where we have used version 4.0 for independent validation and consistency. For completeness in Supplementary Figs. 5C and 6C we report the results obtained with NetMHCpan4.1, showing an improvement of performance for rare alleles, thanks to the increase of training data. These results support our main conclusion that our approach is especially useful in scenarios where new data become available but current methods are not updated yet to cover the corresponding alleles with good accuracy, as was the case when data from Refs. [6, 24] came out while NetMHCpan4.0 was the latest version.

RBM-MHC can also be applied to unannotated, moderate-size multi-allelic datasets available from clinical studies, for which there may exist only a limited number of HLA annotations in existing databases (< 100) for the patient’s rarer HLA alleles. We have shown that RBM-MHC efficiently exploits the statistical information in these samples and combines it with limited annotation information to deliver accurate and stable predictions of HLA-I binding (Figs. 2C-E, Supplementary Figs. 7 and 8, Supplementary Tables 2 and 3).

The flexibility of choice of the training dataset and corresponding applications is even broader. The method is not limited to MS datasets but can be trained on datasets from binding affinity assays to customize presentation score predictions towards the identification of high-affinity ligands (see Fig. 1F). If only MS data are available to build a predictor for this task, as is the case for HLA-C alleles in Fig. 1F, a commonly acknowledged issue is that biases of MS techniques can potentially affect the amino acid frequency distribution in training datasets and hence the predictor’s results. To address this issue we developed, similarly to [11], a sequence re-weighting scheme to efficiently compensate for detection biases in MS and better score ligands tested in *vitro* (STAR Methods, Supplementary Fig. 4).

Overall, our results show that the approach is useful for systematic applications with newly produced large-scale datasets covering an increasing range of HLA types. Future work will be devoted to also develop an extension to HLA class II presentation. Finally, since our method is designed to be trained on custom samples, it could be of relevance to produce sample-specific insights about the complexity of endogenous antigen presentation. As future direction, we will investigate how the probabilistic framework provided by RBM-MHC can be exploited to develop these insights in a quantitative manner.

## Author Contributions

B.B., S.C., R.M., T.M., A.M.W. designed research; B.B., S.C., R.M., T.M., A.M.W. performed research; B.B., J.T., S.C., R.M., T.M., A.M.W. contributed analytic tools; B.B. analyzed data; and B.B., S.C., R.M., T.M., A.M.W. wrote the paper.

## Acknowledgment

We thank Benjamin Greenbaum for suggesting to look at SARS-CoV-2, and helpful exchanges with Clément Roussel, Andrea Di Gioacchino, Carlos Olivarez, Hannah Carter, David G Feller, Shelly Kalaora, Yardena Samuels, David Hoyos, Jayon Lihm, Anna Paola Muntoni. This work was partially supported by Stand Up to Cancer, the European Research Council Consolidator Grant n. 724208, the ANR-17 RBMPro and ANR-19 Decrypted CE30-0021-01, the ANR Flash Covid 19 - FRM, PROJET “SARS-Cov-2immunRNAs”, European Research Council Marie Curie-Sklodowska ITN QuanTI grants. J.T. was supported by a postdoctoral fellowship from the Human Frontier Science Program Organization (reference number: LT001058/2019).

## STAR Methods

### KEY RESOURCES TABLE

| REAGENT or RESOURCE | SOURCE | IDENTIFIER |
| --- | --- | --- |
| Deposited Data |  |  |
| IEDB MHC ligand data | Vita et al., 2019 | http://www.iedb.org/doc/mhc_ligand_full.zip |
| IEDB-derived multi-allelic benchmark samples | This paper | Supplementary Table 2; https://github.com/bravib/rbm-mhc/tree/master/data/Multiallelic_Synthetic_individuals |
| Software and Algorithms |  |  |
| RBM-MHC | This paper | https://github.com/bravib/rbm-mhc |
| NetMHCpan4.0 | Jurtz et al., 2017 | http://www.cbs.dtu.dk/services/NetMHCpan-4.0 |
| NetMHCpan4.1 | Reynisson et al., 2020 | http://www.cbs.dtu.dk/services/NetMHCpan-4.1/ |
| NetMHC4.0 | Andreata et al., 2016 | http://www.cbs.dtu.dk/services/NetMHC |
| MixMHCp2.1 | Bassani-Sternberg and Gfeller, 2016; Gfeller et al., 2018 | https://github.com/GfellerLab/MixMHCp |
| Other |  |  |
| MS-identified peptides in mono-allelic cell lines | Sarkizova et al., 2020 | https://github.com/bravib/rbm-mhc/tree/master/data/Allele-specific_models |
| MS-identified peptides carrying mutations in cancer cell lines SKOV3, A2780, OV90, HeLa, A375 | Marty et al., 2017 | https://github.com/bravib/rbm-mhc/tree/master/data/Allele-specific_models |
| SARS-CoV-2 epitopes (tested *in vitro*) | Immunitrack and Intavis | https://www.immunitrack.com/free-coronavirus-report-for-download/ |
| SARS-CoV and SARS-CoV-2 homolog epitopes | Grifoni et al., 2020 |  |
| Multi-allelic MS cancer sample 12T | Kalaora et al., 2016; Kalaora et al., 2018 | https://github.com/bravib/rbm-mhc/tree/master/data/Multiallelic_Cancer_samples |
| Multi-allelic MS cancer samples Mel8, Mel15, Mel5 | Bassani-Sternberg et al., 2016 | https://github.com/bravib/rbm-mhc/tree/master/data/Multiallelic_Cancer_samples |
| Cosmic Cell Lines Project | Forbes et al., 2016 | https://cancer.sanger.ac.uk/cosmic |
| Genomics of Drug Sensitivity Center | Yang et al., 2013 | https://www.cancerrxgene.org/ |
| Single Nucleotide Polymorphism Database (dbSNP) | Sherry et al., 2001 | http://www.ncbi.nlm.nih.gov/SNP/ |
| GenBank | Benson et al., 2013 | https://www.ncbi.nlm.nih.gov/genbank/ |
| UniProt | The UniProt Consortium 2019 | http://www.uniprot.org/ |
| HLA frequency net | González-Galarza et al., 2015 | http://www.allelefrequencies.net/ |

## RESOURCE AVAILABILITY

### Lead Contact

Further information and requests for resources and reagents should be directed to and will be fulfilled by the Lead Contact, Barbara Bravi.

### Materials Availability

This study did not generate new materials.

### Data and Code Availability

The RBM-MHC source code, datasets and trained models supporting the current study are available at https://github.com/bravib/rbm-mhc.

## METHOD DETAILS

### Schematic outline of the RBM-MHC pipeline

The essential steps of the RBM-MHC application pipeline are:

1. *Collection of training datasets*. RBM-MHC can be trained on patient-derived clinical samples or on datasets collected from public repositories - here we constructed training datasets from mass spectrometry and binding affinity assays available in IEDB. See Supplementary Fig. 1A and STAR Methods section “Dataset collection from IEDB and preparation” for a detailed description.
2. *Alignment*. RBM-MHC works with fixed-length sequences, hence peptide sequences are first reduced to the same length through an alignment routine, see Supplementary Fig. 1E and STAR Methods section “Antigen sequence alignment” for a detailed description.
3. *RBM-MHC algorithm*. The full RBM-MHC architecture (Fig. 2A) combines a Restricted Boltzmann Machine (RBM) and a HLA-I classifier, trained in successive steps. See Supplementary Figs. 2, 3 and STAR Methods section “RBM-MHC algorithm” for a detailed description.

### Dataset collection from IEDB and preparation

The analyses described throughout this work rely on the collection of sets of peptides documented with selected HLA restriction in the Immune Epitope Database (IEDB, last release [13]).

#### Mass spectrometry datasets

The search in IEDB for immunopeptidomic data from mass spectrometry (MS) was performed as follows. The full set of curated HLA-I ligands was downloaded from IEDB (file *mhc_ligand_full.csv* from http://www.iedb.org/database_export_v3.php, accessed in July 2020). In this IEDB file, we looked for linear, human peptides eluted from cells and detected by MS techniques (field *Assay Group* = “ligand presentation”, field *Method/Technique* = “cellular MHC/mass spectrometry”, “mass spectrometry”, “secreted MHC/mass spectrometry”). Among these data, we prioritized HLA-specific peptides from mono-allelic sources, in such a way that the assignment of HLA-binding specificity is unambiguous and does not rely on additional *in silico* predictions. To do this in practice, we set the field *Allele Evidence Code* = “Single allele present”, indicating that antigen presenting cells are known to only express a single HLA-I molecule, as is the case for mono-allelic cell lines [4, 6]. If, for a given allele, we found less than a minimal number (set to 300) of sequences among mono-allelic-source data, we extended the search to peptides with *Allele Evidence Code* = “Allele Specific Purification”, since this procedure attaches greater confidence to the HLA assignment than its inference by *in silico* methods. Only for one allele (HLA-B*39:01) did we extend the search to all other MS data, including the evidence code “Inferred by motif or alleles present”. MS datasets filtered through these steps were used to train allele-specific RBM presentation models (as in Figs. 1C-F) and multi-allelic models (as in Fig. 2C-E). In a multi-allelic setting, we first tested motif reconstruction by considering 10 “synthetic-individual” datasets of antigens with known HLA-I specificity to assess the RBM-MHC classification performance by comparing RBM-MHC predictions against the known HLA-I specificity. These datasets were built by collecting all IEDB antigens (searched as above) associated to 2 haplotypes, *i.e.,* combinations of an A, a B, a C HLA allele observed to co-occur in the human population to preserve linkage (see Supplementary Table 2). Information on haplotypes co-occurrence was found at allelefrequencies.net [34]. To apply the RBM-MHC method to multi-allelic, patient-derived immunopeptidomic samples, we sought to have a small amount of labeled peptides, here set to 10%. To this end, we either retrieved a HLA annotation in IEDB for portions of the samples or, when this was not possible, we added to each of them labeled peptides from IEDB (searched as above) for the 6 HLA-I given by the patient’s HLA typing. The RBM-MHC training set was then defined as this minimally extended dataset, to guide the learning of the HLA-I classifier by the labeled peptides, whose predictions are used to reconstruct HLA-I motifs in the original dataset. We did not attempt to identify motifs in the dataset Mel5 from [31] as the patient’s HLA typing was incomplete (lacking the HLA-C alleles).

#### Binding assay datasets

Records of binding affinity (BA) assays from IEDB (as of July 2020) were filtered following [6], *i.e*. selecting peptides annotated with a quantitative measure of binding dissociation constants < 500 nM and excluding assay types that led to documented discrepancies between predicted and effective affinity (“purified MHC/direct/radioactivity/dissociation constant KD”, “purified MHC/direct/fluorescence/half maximal effective concentration (EC50)”, “cellular MHC/direct/fluorescence/ half maximal effective concentration (EC50)”). BA datasets were used to train allele-specific models of Figs. 1F-G.

#### Antigen sequence alignment

The typical length of HLA-I peptides, structurally constrained by the MHC-I binding cleft, is generally recognized to be 8 to 11 amino acids [6, 7, 17, 18], with 9-mers being the most abundant except for very few exceptions [6]. Hence we focus on datasets containing 8-11-mers (Supplementary Figs. 1B-D) and an alignment procedure is implemented to reduce peptide sequences to the typical length of 9 residues. This choice of a 9-mer-centered alignment is also consistent with the treatment of variable-length class I peptides by other algorithms as MixMHCp, which scores their positions based on 9-mer motifs [12], and NetMHC, which applies to sequences with length different from 9 the insertions and deletions that give optimal predicted scores for presentation by a given allele [7].

Our alignment routine is articulated around the construction of a main alignment and an alignment refinement, aimed at improving HLA classification and hence motif reconstruction. The optimal alignment for each peptide is found by separately aligning subsets of peptides sharing the same HLA-binding specificity that best describe the corresponding sequence motif. In turn, optimally aligning a peptide facilitates its correct association the HLA type. However, identifying such subsets in typical samples, which pool together peptides of different specificity, requires first a step of motif reconstruction, *i.e*. of assignment of putative HLA types to all peptides. The workflow of the alignment procedure, depicted in Supplementary Fig. 1E, is as follows.

Main alignment:

- Progressively align fixed-length subsets of sequences (Step 1 in Supplementary Fig. 1E). We estimate Position Weight Matrix (PWM) profiles of subsets of sequences with the same length and we align these profiles, using respectively the functions *seqprofile* and *profalign* (with default options) of the Matlab Bioinformatics Toolbox (release R2018b). The alignment is made progressively from the minimal length considered (here 8 residues) to the maximal one (here 11 residues) by inserting gaps in shorter profiles, resulting in an alignment of the maximal length considered (11). This alignment is used as seed to learn a Hidden Markov Model (HMM) profile of the reference length = 9 by appeal to the routines *hmmprofstruct* and *hmmprofestimate* (with default parameters) of the Matlab Bioinformatics Toolbox.
- Align sequences to the HMM profile, Step 2 in Supplementary Fig. 1E. Sequences with length different from 9 are re-aligned to the HMM profile relying on the position-specific insertion/deletion probabilities of the HMM (by the *hmmprofalign* function). This procedure results in a multiple sequence alignment displaying mostly the insertion of a gap in 8-mers and single/double deletions in 10/11-mers.
- Use this first alignment to train RBM-MHC.

Alignment refinement (for HLA classification in multi-allelic samples):

- Build HLA-specific HMM profiles by grouping peptides based on the putative “class” (HLA type) predicted by the HLA-I classifier (Step 3 in Supplementary Fig. 1E). First, for each HLA-I class, we put together the 10% of labeled data and the peptides classified in that class, weighted by the probability of classification, reflecting the degree of confidence of class assignment. We build on this sample a HLA-specific HMM by the routines *hmmprofstruct* and *hmmprofestimate* (with default parameters). We use these HMM profiles (essentially capturing the pattern of single-site amino acid frequency for each HLA type) as seeds of each class’s alignment.
- Re-align peptides based on the best HMM alignment scores (Step 4 in Supplementary Fig. 1E). We take every unlabeled peptide and we consider the alignment to each of the classes’ seeds and the corresponding HMM alignment score (both given by the *hmmprofalign* function). We retain, for each peptide, the alignment with the highest score. Such best scoring alignment can be to a class that is different from the classifier-predicted one, allowing us to re-classify peptides more accurately in the subsequent iteration. This step helps therefore correct classification errors arising from the initial, suboptimal alignment by means of allele-specific HMM alignment scores.
- Repeat the RBM-MHC training after the re-alignment.

In the 10 “synthetic-individual” datasets considered for testing motif reconstruction (Supplementary Table 2), high classification performance was reached already at the first iteration and we observed a further, systematic improvement after 1 re-alignment step, see Supplementary Fig. 1F. For 2 or more iterations, there are cases in which the classification performance is degraded, therefore in our motif reconstruction applications (Figs. 2C-E) we re-align and re-train once.

The alignment routine is designed in such a way as to be easily tuned to a different reference length and to the inclusion of longer peptides, depending on the particular length preferences of the alleles of interest and on data availability. For instance, in Supplementary Fig. 1H we show that HLA classification performance is rather stable with respect to the choice of the reference length in the range 8-11 residues and could even slightly increase for reference lengths > 9. We recall however that the choice of reference length = 9 is made to reflect the typical length of class I peptides: peptide length distributions of the datasets considered in this work confirm that 9 residues is by far the most abundant length (Supplementary Figs. 1B-D).

To test the quality our alignment routine, we compared its performance at HLA classification to that of other common alignment software, MAFFT (latest version 7.471 [29]) and HMMER (http://hmmer.org/). Default options of these methods tend to produce very gapped alignments, that might cause a rather drastic loss of information on the original peptides and hence affect the RBM-MHC classification performance. In contrast, the progressive alignment of profiles obtained from subsets of sequences of the same length (Step 1 in Supplementary Fig. 1E) allows for well controled gap insertions. We found that large gap penalties are more suitable for applying MAFFT to the type of alignment of interest here, and yield better performance. After a grid search we set in MAFFT the penalty for opening a gap to 8, the one for extending a gap to 12. We implemented MAFFT: *(i)* with a speed-oriented option, FFT-NS-2, see [29]; *(ii)* with an accuracy-oriented option - G-INS-i - recommended for sequences of similar lengths. (We have checked that other accuracy-oriented options, E-INS-i and L-INS-i, lead to rather similar results). MAFFT accuracy-oriented options tend to be slow, hence we ran it only on a subset of the sample. We use this aligned subset as a seed to build a HMM by the HMMER routine *hmmbuild* (with open-gap penalty and extension-gap penalty respectively increased to 0.49 and 0.99). All the other sequences are then aligned by the routine *hmmalign*. To get an alignment of length 9, one can adjust accordingly a maximal gap percentage, i.e., columns with more than this percentage of gaps are filtered out to reduce the alignment length. For example, to obtain the alignments of length = 9 used for Supplementary Fig. 1G, columns with in average > 68% of gaps (MAFFT-FFT-NS-2) and > 65% of gaps (MAFFT-G-INS-i + HMMER) were filtered out. Varying this percentage threshold allows one to work with alignments of different reference length, see *e.g*. Supplementary Figs. 1I-L. We did not find evidence that MAFFT-based alignments *(i)* and *(ii)* lead to improved performance compared to the alignment routine we adopted, already in the first training iteration (Supplementary Fig. 1G). In particular, the fast, but usually less accurate, FFT-NS-2 option, allowing for a global alignment of the full sample, performs better than the use of a HMM model built from an accurately aligned seed. This might be due to the fact that the initial choice of a unique seed, including peptides of different specificity yet to predict, is inevitably suboptimal.

### RBM-MHC algorithm

#### Restricted Boltzmann Machine (RBM)

The RBM learns an underlying probability model for the antigen sequences in the training datasets, in our case the HLA-I presented antigens. A RBM [14, 15] consists of one layer of *N^v^* observed units 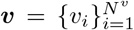 and one layer of *N^h^* hidden units 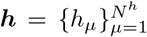 connected by weights ***W*** = {*w_i_μ__*} (see Supplementary Fig. 2A). The observed units ***v*** stand here for the antigen sequences to model, hence *N^v^* = 9 and the number of “symbols”, *q*, for each observed unit is 21 (20 amino acids and the gap). Mathematically, the model is defined by a joint probability over presented antigen sequences and hidden units:

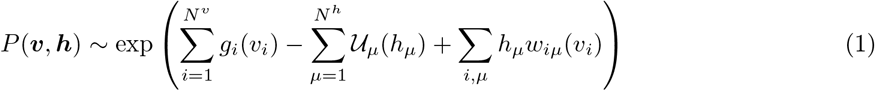

where *g_i_*(*v_i_*) is a matrix of *N^v^ × q* local “potentials” (biases) acting on observed units, 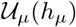 are *N^h^* local potentials on hidden units and the weights *w_i_μ__*(*v_i_*) (arranged in a tensor of size *N^v^ × N^h^ × q*) couple hidden and observed units. The parametric form of 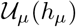 is chosen as a double Rectified Linear Units (dReLu):

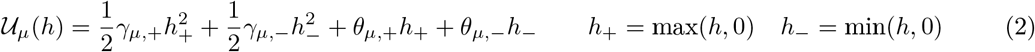

containing parameters (*γ_μ_,+, γ_μ,−_, h_+_, h_−_*) to infer from data during training (see below). The dReLu was shown to outperform other choices of potential such as Gaussian [16, 35], guaranteeing that correlations beyond pairwise in data are captured. The probability distribution over the presented antigen sequences one is interested in modeling is recovered as the marginal probability over hidden units:

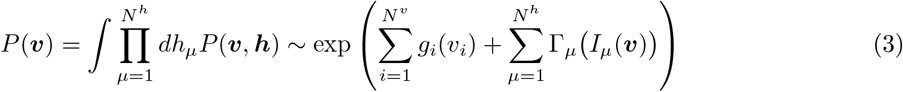

where 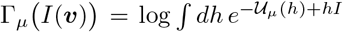. We define *I_μ_*(***v***), the input to hidden unit *μ* coming from the observed sequence ***v***, as the sum of the weights entering that particular hidden unit:

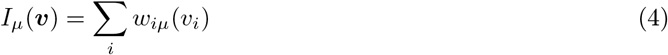

During the modeling step called “training”, the weights *w_i_μ__*, the local potentials *g_i_*(*v_i_*) and the parameters specifying 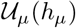 are inferred from data by maximizing the average log-likelihood 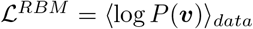 of sequence data ***v***, as previously described [16, 35]. This leads to inferring the RBM probability distribution that optimally reproduces the statistics of the training dataset (see Supplementary Fig. 3H for comparisons of data and model single-site frequency and pairwise connected correlations). During training, the contribution of sequences to 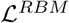 can be re-weighted following a re-weigthing scheme we have conceived (see STAR Methods section “Sequence re-weighting scheme”) to correct for amino-acid frequency biases in the training dataset, as the ones introduced by mass spectrometry (MS) detection. A regularization, *i.e*. a penalty term over the weight parameters is introduced to prevent overfitting. The function maximized during training then becomes:

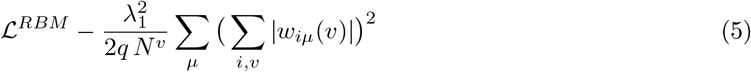

This regularization, which was first introduced in [16], plays effectively the role of a *L*_1_ regularization, imposing sparsity of weights, with a strength that is adapted to increasing magnitude of weights, hence favouring homogeneity among hidden units (see [16] for more detailed explanations). Examples of inferred sets of weights at different regularization strength 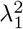 are provided in Supplementary Figs. 2B and 3G. The package used for training, evaluating and visualising RBMs is an updated version of the one described in [16]. The package was ported to Python3 and the optimization algorithm was changed from SGD to RMSprop (*i.e*., to ADAM without momentum) with learning rate 5 · 10^−3^, *β*_1_ =0, *β*_2_ = 0.99, *ϵ* = 10^−3^) (see [36] for definition of the parameters). The parameter values were chosen for their robust learning performance across a variety of datasets studied in previous works such as MNIST, Ising models, and MSAs of protein domains of various sizes. Overall, the adaptive learning rate of RMSprop/ADAM result in larger updates for the fields and weights attached to rare amino acids, and hence speeds up convergence.

The (hyper-)parametric search for optimal regularization 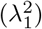 and number of hidden units (*N^h^*) was made from the trend of the RBM log-likelihood on a held-out validation dataset, the aim being to achieve a good fit but to avoid overfitting. Supplementary Figs. 3A-B illustrate such (hyper-)parametric search for a multi-allelic RBM. Low regularizations achieve a better fit of training data (see log-likelihood values for the training dataset); when selecting a low regularization (as 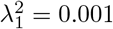), the log-likelihood over the validation dataset starts to decrease beyond *N^h^* = 10, indicating overfitting. Given these trends, we trained the multi-allelic RBM models (Figs. 2C-E) with *N^h^* = 10 and 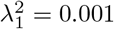. This choice is further supported by considering, in Supplementary Fig. 3C, the accuracy of HLA classification (see below for its definition), which reaches an optimal value for 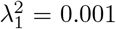 and *N^h^* = 10. The same test with an allelespecific RBM (Supplementary Figs. 3F) shows that already beyond *N^h^* = 5 the model could overfit, hence allele-specific RBM presentation models (Figs. 1C-G) were trained with *N^h^* = 5 and 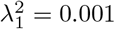.

#### HLA-I Classifier

The HLA-I classifier part of the RBM-MHC takes as input 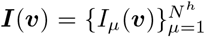 of the peptide sequence ***v*** and gives a categorical output, *i.e*. the peptide HLA-I specificity (Fig. 2A and Supplementary Fig. 2A). *c* = 1,…, *N^c^* denotes the HLA-I type. Typically for single-individual samples *N^c^* = 6, since each individual displays at most 2 different copies (a maternal and a paternal copy) of HLA-A, HLA-B and HLA-C.

The classifier is trained by minimizing a loss function chosen to be a categorical cross-entropy 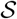:

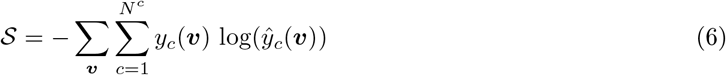

where the ***v*** sum runs only over the sequences labeled with their HLA-I association. *y_c_*(***v***) is the label assigned to ***v*** for supervised training in one-hot encoding, *i.e. y_c_*(***v***) = 1 only for the c standing for its associated HLA type and zero otherwise so ∑*_c_y_c_*(***v***) = 1 (it is normalized over categories). The choice of one-hot encoding is justified by the fact that, for the sake of discriminating motifs, we select peptides associated to only 1 HLA type in the database (mainly from mono-allelic sources, see STAR Methods section “Dataset collection from IEDB and preparation”). 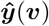 is the categorical output predicted by the HLA-I classifier for ***v***, calculated from the softmax activation function:

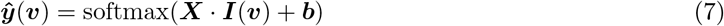

where ***X*** are the classifier weights, connecting input to output layer (see Supplementary Fig. 2A), and ***b*** are local biases adjusted during training. Element-wise softmax is defined as:

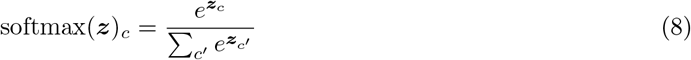

The activation function softmax has the advantage of giving predictions normalized over categories, thus each element 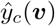 can be interpreted as the probability that sequence ***v*** belongs to class *c*. Our numerical implementation relies on Theano and Keras Python libraries. Training is performed in minibatches of 64 sequences, by the ADAM optimizer for 1000 epochs, retaining the model that gives the best accuracy on a held-out partition of the ~30% of the training dataset. The choice of RBM (hyper-)parameters (see above) also ensures a high accuracy of classification (Supplementary Fig. 3C). Accuracy of classification is measured there as an Area Under the Curve (AUC), which is different from the cross-entropy optimized during training (Eq. 6) and is defined below, in the section “RBM-MHC performance in multi-allelic samples”. This AUC value is only minimally affected when reducing the RBM training dataset and is close to 1 (indicating the maximal accuracy) already with the small number of labeled sequences used, *i.e*. 10% of data. (The AUC clearly increases further when increasing this amount, see Supplementary Fig. 3D-E).

Combining the probability functions estimated by RBM and HLA-I classifier, we define for each sequence ***v*** a global score 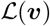:

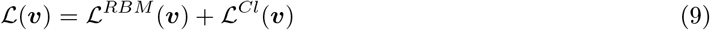

where 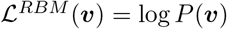 is the RBM log-likelihood assigned to sequence ***v*** and 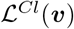 is the classifier score, defined from the vector of predicted class probabilities 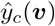 as 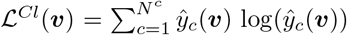. 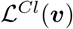 is the negative entropy of classification, so it contributes to 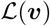 with higher values when the confidence with which a HLA-I class is predicted is higher.

We shall stress here that we do not attempt to optimize log-likelihood given by the global score Eq. 9, as this strategy has the risk of introducing biases in the representation learnt by the RBM in a fully unsupervised way - backpropagating the classifier gradient to the RBM training would have the effect of adapting its hidden-space representation to the prediction of the HLA type. Rather, it is precisely learning the classifier on top of the cluster structure discovered in an unsupervised way in the RBM representation space that ensures robust HLA type prediction in the scenario we envision, *i.e*., when we have very few labels at our disposal to characterize new datasets. In addition, optimizing Eq. 9 could potentially also imply a loss of the feature discovery power of a fully unsupervised approach, as data features not directly contributing to the HLA specificity would not be internally represented.

### Unsupervised clustering

#### K-means algorithm

Given a set of points ***x***, K-means [28, 37] finds the centroids of *c* = 1,…, *N^c^* clusters and assigns each ***x*** to the cluster whose centroid has the minimal distance to ***x***. If we indicate by *d_c_*(***x***) the distance between point ***x*** and a cluster c, we can express the probability that ***x*** belongs to cluster *c* as:

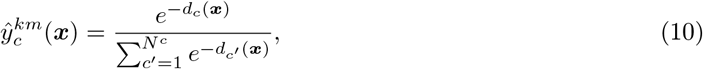

as is a common choice in the “soft” version of K-means. We define RBM-km as the application of K-means to sequence representations in the space of inputs to hidden units (*i.e. **x** = **I***(***v***)) instead of sequences themselves (*i.e. **x** = **v***). From such a probabilistic clustering prediction, we define the classification score 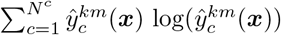. Our implementation of K-means relies on the routine available in the Python package Scikit-learn [38].

#### MixMHCp algorithm

Consistently with the other approaches discussed, we assume that the expected number of clusters *N_c_* is known, and we implement the clustering by MixMHCp2.1 [10–12]. First we build Position Weight Matrices (PWM) for each of the *N^c^* clusters found by MixMHCp; each PWM describes the single-site amino acid frequencies 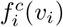 in cluster *c*, with *c* = 1,…, *N^c^*. For a sequence ***v*** a set of presentation scores (one per cluster) can be defined from the log-likelihood under the corresponding PWM:

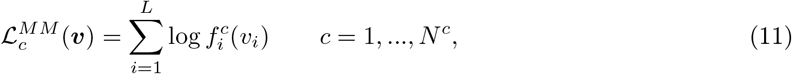

where the superscript *MM* stands for “MixMHCp” and *L* is the length to which all sequences are reduced in MixMHCp (*L* = 9). The final score of a sequence ***v*** is taken as the maximal log-likelihood among the *N^c^* clusters, 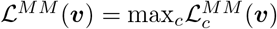, which is used jointly to predict the HLA association.

#### Annotation of HLA-I binding motifs

The first step in clustering approaches, either by MixMHCp or K-means, is fully unsupervised and consists of optimally assigning peptides to *N^c^* clusters. Clusters then need to be annotated with the corresponding HLA-binding specificity, among the *N^c^* ones known to be expressed in the sample cells from *e.g*. HLA typing. For this second step, we consider the fraction of data for which we have labels (the same used for training the HLA-I classifier) and we estimate from them a PWM for each HLA type, in such a way as to obtain a set of reference motifs. We next estimate the PWM of each cluster and we label the cluster with the HLA association of the reference motif that minimizes the difference to the cluster PWM. Note that this mapping could give the same HLA type associated to several clusters and other HLA types not associated to any cluster, indicating a poor classification performance.

### Generation of HLA-specific artificial peptides

A RBM is a generative model. As it is based on fitting an entire probability distribution to a given dataset, sampling from this distribution allows one to generate new candidate antigens. The binding specificity to HLA-I of such generated sequences can be controlled by conditioning (fixing) the RBM probability on the values of inputs to hidden units coding for the desired specificity. The search for these values is guided by the HLA-I classifier. This procedure directly builds on the idea of sampling while conditioning on structural, functional, phylogenetic features emerging in RBM representations of protein families [16]. Schematically, the steps of this HLA-specific sampling are as follows:

1. we select a HLA-I class *c* (e.g. *c* = HLA-A*01:01) and we find ***I**^o^* such that 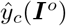 is close to 1 (that is, there is high confidence of class *c* prediction);
2. we estimate ***h**^o^* = 〈***h***〉 from the conditional probability *P*(***h**|**I**^o^*) and we sample new sequences ***v*** from the probability of observed sequences conditioned on ***h**^o^, P*(***v**|**h** = **h**^o^*);
3. to further explore the region encoding for the specificity to HLA-I class *c*, we randomly move from ***I**^o^* to ***I**^n^ = δ**I**^o^ + δ**I*** (*δ**I***s drawn from a Gaussian 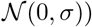);
4. we accept the move with a probability *π* ~ exp(−*β*(*e^n^ − e^o^*)), where 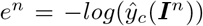 and 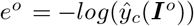. The parameter *β* (akin to an inverse temperature in physics) basically controls how stringent the selection for sequences predicted in class c with a high probability is.
5. We set ***I**^o^ = **I**^n^* to proceed with new moves as in step 3. Every arbitrary amount of moves, we generate new configurations by conditional sampling as in step 2.

Supplementary Fig. 2C shows examples of ***I**^n^* values (2 of its components for simplicity) covered in this search with *β* = 50 and *σ* = 0.01 (dark red points). By this sampling procedure, we generate samples of artificial peptides that would be predicted to be presentable and specifically presented by the selected HLA-I protein complex. Such samples explore a broad diversity of peptides, that are typically 3-4 mutations away from the closest natural ones (Supplementary Fig. 2D) while preserving the profile of amino acid abundances constrained by a given binding specificity (Supplementary Fig. 2E).

### Sequence re-weighting scheme

RBM presentation models trained on datasets obtained by mass spectrometry (MS) might underestimate the probability of presentation of ligands that are less frequently detected by MS. The most evident case are ligands containing cysteine, an amino acid that can undergo chemical modifications by oxidation typically not included in standard MS searches. We introduced a procedure for re-weighting sequences as a general strategy to correct for biases in the training dataset, such as MS detection biases. We defined a weight for each sequence ***v*** in the training set:

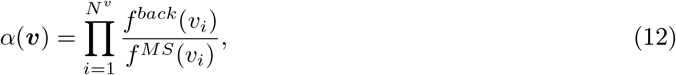

where *f^MS^* denotes typical amino acid frequencies in MS (as can be estimated from IEDB data) while *f^back^* is a background amino acid frequency distribution that does not suffer from the same biases as the MS one. These weights *α*(***v***) are incorporated into the average log-likelihood to maximize during RBM training:

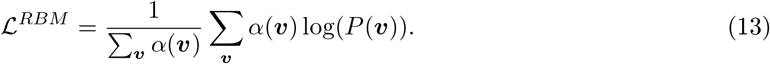

When modeling multiple specificities through mixtures of PWMs (as by MixMHCp and MixMHCpred, see [11]), an analogous correction can be implemented by rescaling the amino acid frequency composing each PWM by the background frequency. Following Ref. [11], *f^back^* can be chosen as the frequency of amino acids in the human proteome or as the frequency of amino acids in IEDB antigens detected by other techniques than MS (we shall refer to these as *f^non–MS^*). More precisely, *f^non−MS^* is estimated from frequencies at non-anchor sites (position 4 to position 7 of 9-mers), excluding from the search all alleles that: *(i)* have fewer than 100 peptides associated to them; *(ii)* are present in neither MS nor non–MS datasets; *(iii)* show specificity at positions between 4 and 7.

In Supplementary Fig. 4A, the comparison of *f^non−MS^* and *f^MS^* for the 20 amino acids provides a clear indication of the MS detection bias in relation to the amino acids cysteine (C) and tryptophan (W), the frequency difference between MS detection and other techniques being more than 100% of MS frequency itself. The C/W frequency in presented antigens is therefore expected to be underestimated by MS, suggesting that ligands containing C/W, whose binding affinity to HLA-I could be successfully tested *in vitro*, would be missed by MS. For the purpose of illustration, we test how the re-weighting procedure could correct for such bias on a presentation model built from IEDB, MS data for the allele HLA-A*01:01, with *f^back^ = f^non−MS^*. We first compute the weights *α* (Eq. 12) for sequences in the training dataset. In Supplementary Fig. 4B, we show separately their distribution for sequences with and without C/W: for the former, *α* have particularly high values, in such a way as to weight more their statistical information and to compensate for the underestimation of C/W frequencies in MS. The effect of re-weighting can be then assessed on a validation dataset, in particular on its sequences with C/W, looking at the distribution of their presentation scores relative to the average scores over the full validation set (see Supplementary Fig. 4C). When considering a RBM model trained without re-weighting (giving the probability *P^MS^*), these sequences would be assigned a score lower than the average score of the validation set, as a straight consequence of the low amount of ligands with C/W in the training set. Once the RBM model is learned with re-weighting (giving the probability 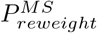), the score assigned to these sequences is brought closer to the average value and in particular to overlap with the range of presentation score values that would be assigned by a RBM model (*P^non−MS^*) trained on IEDB data for the same HLA allele from techniques different from MS (*i.e*. data that do not underestimate the occurrence of C/W in presented antigens). The re-weighting scheme can also improve the ability of the RBM model to discriminate antigens containing C/W of validated immunogenicity from generic peptides, by assigning them presentation scores of higher rank. We show this in Supplementary Fig. 4D, where we consider a re-weighted version of the HLA-A*02:01-specific model of Figs. 1C,E. While there (as well as in the analysis of Figs. 1D and 2B-E) the re-weighting was not necessary, as we validated the model on the same type of data (MS-detected) as the training set (MS-based), in Supplementary Fig. 4D we show that the re-weigthing is useful for scoring by a MS-based model peptides from binding assays and that contain amino acids underdetected by MS. Similarly, another scenario where we applied the re-weighting scheme is the prediction, by MS models, of SARS-CoV-2 epitopes tested *in vitro* for binding, as in Fig. 1F. The re-weighting procedure just described can be activated as an option in the RBM-MHC software package.

## QUANTIFICATION AND STATISTICAL ANALYSIS

### Model predictions in mono-allelic MS datasets

To assign scores of presentation by a specific HLA-I, we resorted simply to the RBM to build HLA-specific presentation models (with N^h^ = 5, 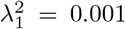). To train them, sets of 8-11-mer peptides documented with the corresponding HLA association were collected from IEDB, restricting to mono-allelic-source, mass spectrometry sequence data, as outlined in the section “Dataset collection from IEDB and preparation”. We then used the RBM log-likelihood 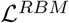 as probabilistic score of presentation by the HLA under consideration. For the preliminary validity check of the HLA-A*02:01-specific model, we randomly selected a subset with the 80% of these sequences as training dataset and we kept the remaining 20% as test set to evaluate the model’s predictions in terms of probabilistic scores of presentation (Fig. 1C). We randomly selected 5000 peptides from the human proteome (as in UniProt database [39]) with length distribution matching the one of presented peptides to serve as a set of peptides predominantly not presented on the cell surface (“generic”).

To test our method on mono-allelic MS datasets from Ref. [6], we trained 92 allele-specific RBM models, for all the A, B, C HLAs analyzed in [6]. We decided to train a series of allele-specific models to minimize the uneven accuracy across alleles, emerging especially in multi-allelic models, due to the different abundances of peptides with different HLA-I preferences available as training data. (The use of multi-allelic models is intended as tool to characterize unannotated samples through motif reconstruction). For each model, we randomly selected in the corresponding mono-allelic dataset 100 peptides to use for model evaluation. Since datasets from [6] feature in the latest version of IEDB, we carefully excluded the 100 peptides per model from the IEDB-derived training datasets. We produced, as described above, *n*-fold excess of generic peptides, choosing *n* in such a way as to consider a proportion between presented and generic peptides that resembles natural conditions of epitope selection. Large-scale experimental and bioinformatic studies on viral peptidomes [19–21] found that ~ 1% of peptides bind to given MHCs (suggesting the value n=99). Comparing the amount of all 9-long peptides from human proteome (~ 10^7^) and the average, allele-specific antigen repertoire size (~ 10^4^ sequences) derived from existing databases as IEDB, Refs. [4, 6] concluded rather that ~ 0.1% of peptides is “presentable” (corresponding to n=999). These approximate estimates allowed us to fix a threshold for positive prediction at, respectively, the top-scoring 1% and 0.1% of peptides in order to evaluate model performance by the PPV metric. (The random expectation for the PPV in these conditions is 10^−2^ and 10^−3^ for the PPV estimated respectively at the top scoring 1% and 0.1%). AUCs do not vary, apart from noise, with *n*; the AUC values shown in Fig. 1D and Supplementary Fig. 5 are the ones obtained with *n*=99 (i.e., generic and presented peptides mixed in proportion 100:1). To better rationalize differences between RBM and NetMHCpan4.0 in connection to different HLAs, we monitored performance as a function of the distance between the query HLA and its nearest neighbor in the NetMHCpan4.0 training dataset, where such distance is determined from the similarity between the two HLA-I sequences (more precisely, the 34 residues in contact with the peptide). The distance between sequences *X* and *Y* is defined as 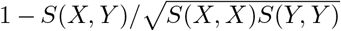, where *e.g*. *S*(*X, Y*) is a global alignment score between *X* and *Y* based on the BLOSUM50 substitution matrix, see [22, 40].

### MS-based model validation for neoantigen discovery

To evaluate the predictive power of the HLA-A*02:01-specific RBM model for neoantigen discovery, we acquired the list of missense mutations for ovarian and melanoma cancer cell lines SKOV3, A2780, OV90, HeLa and A375 from the Cosmic Cell Lines Project [41]. We further filtered them largely following the steps described in Ref. [23], *i.e*. we retained only mutations in genes with non-zero expression levels in the corresponding cell line (based on expression data from the Genomics of Drug Sensitivity Center [42]) and we excluded common germline variants documented in the Single Nucleotide Polymorphism Database [43].

As a result, the number of missense mutations considered is as follows: 444 (A375), 521 (SKOV3), 602 (A2780), 280 (OV90), 511 (HeLa). For each cell line, the list of mutated peptides to score results from adding up all the up to 38 peptides of length 8-11 containing each mutation. Since the MS database for the 5 cancer lines produced by Ref. [23] was annotated by comparison to the consensus human proteome, it contained only wildtype peptides whose mutated version was in the list of putative neoantigens. As long as a mutation does not affect an anchor site, the mutated version should preserve a high probability of presentation. We followed the authors’ choice of not considering the two peptides mutated at anchor positions as presented and we excluded them from the analysis of Fig. 1E. When we consider all neoantigens (including the two neoantigens arising from anchor-site mutations of MS-derived peptides) the mean score percentiles are: RBM-MHC 3.3% *vs* NetMHCpan4.1 1.6% (among generic peptides) and RBM-MHC 3.6% *vs* NetMHCpan4.1 0.9% (among all mutated peptides from the same cell line).

The studies from which single-patient samples 12T, Mel8, Mel15, Mel5 were retrieved [26, 27, 31] listed a total of 11 tumor-presented neoantigens, using techniques that allow for the detection of variants of native proteins by MS. For some of these neoantigens, immunogenicity was also validated *in vitro* by identification of neoantigen-specific T-cell responses. Their HLA association was predicted by NetMHC software, see [26, 27, 31]. The prediction was to a large extent confirmed by *in vitro* validation of immunogenicity, which relied on antigen presenting cells that were positive to the predicted HLA-I. As the 11 neoantigens are 9-10 residues long, following Ref. [11] we considered 9-mer and 10-mer peptides overlapping the missense mutations observed in the patient’s WES, which resulted in a list of thousands of putative neoantigens (see Supplementary Table 3). The RBM-MHC training set *per se* can thus take into account only 9-10 mers from patients’ samples; patients’ neoantigens to validate were not included in the training dataset. In this multi-allelic case where antigens may be presented by several HLA types (as opposed to the allele-specific case of the previous paragraph) we used the global score accounting for the probability of presentation (by RBM) as well as the confidence of the HLA assignment (either by the HLA-I classifier in RBM-MHC or by K-means clustering in RBM-km) of Eq. 9.

### Model predictions for SARS-CoV-2 epitopes

We downloaded the protein-coding regions of SARS-CoV-2 genome from GenBank [44], NCBI Reference Sequence: NC_045512.2. We extracted all the 9-mers contained in the SARS-CoV-2 proteome, giving a list of 9656 candidate cytotoxic T-cell epitopes (HLA-I antigens). Ref. [24] (available at https://www.immunitrack.com/free-coronavirus-report-for-download/) tested *in vitro* the 94 top scoring epitopes according to NetMHC4.0 [7] (a well-known NetMHC version for binding affinity predictions) for each of the HLA-I alleles A*01:01, A*02:01, A*03:01, A*11:01, A*24:02, B*40:01, C*04:01, C*07:01, C*07:02 and according to NetMHCpan4.0 (run with EL option) [8] for C*01:02. 159 peptides were identified as high-stability binders (i.e., with stability above the threshold of 60% of the reference peptide value for the corresponding HLA-I allele). Ref. [24] also estimated the trend of binding stability (expressed as % of the reference value) *vs* predicted score (binding affinity in [nM] for NetMHC4.0, rank percentile for NetMHCpan4.0). Here we report and quantify these trends in terms of Pearson correlation for the sake of comparison to RBM (Fig. 1F and Supplementary Fig. 6B). To probe our method as a predictor for the high-stability binders found, we learned a series of allele-specific RBM presentation models (*N^h^* =5 hidden units, 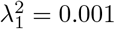) for the 10 HLA-I alleles considered in this study, prioritizing training datasets from binding assays in IEDB (see STAR Methods section “Dataset collection from IEDB and preparation”). These types of data are almost absent for the 4 HLA-C mentioned, and in these cases models were learned from MS data only but a re-weighting of frequencies, aimed at correcting for MS biases in detection, was applied (see STAR Methods section “Sequence re-weighting scheme”). Final BA datasets used have in average 1269 sequences, MS ones 1678. All peptides tested for binding to a given HLA-I in Ref. [24] were scored by the corresponding RBM model (see Fig. 1F, where, as a comparison, results achieved by training all RBMs on MS data with re-weigthing are also reported). We next estimated their score percentile relative to scores assigned to all candidate epitopes and we assessed that tested binders were predominantly assigned high scores, able to a good extent to discriminate them from the tested non-binders (see Supplementary Fig. 6A).

As an additional test, we considered the SARS-CoV-2 cytotoxic T-cell epitopes identified as potentially associated to high immune responses by Ref. [25], who mapped the dominant, experimentally validated SARS-CoV-derived epitopes to the corresponding regions in the SARS-CoV-2 proteome. We scored the 22 epitopes in this list with complete (100%) or moderate-high (> 70%) sequence similarity to the homologous SARS-CoV epitope, which should preserve high likelihood of presentation. Since homologous SARS-CoV epitopes were experimentally tested in binding assays, scores were assigned by the same (affinity-trained) models as above covering the HLA restrictions reported in Ref. [25] (HLA-A*02:01 to the largest extent, HLA-A*24:02, HLA-B*40:01), see Fig. 1G and Supplementary Fig. 6D. In addition, we checked that scores estimated by MS-trained RBM models with re-weighting lie in the same range, see Supplementary Fig. 6E.

### RBM-MHC performance in multi-allelic samples

We chose the Area Under the Curve (AUC) of the Receiving Operating Characteristic (ROC) as a metric for classification (i.e., HLA assignment) performance, estimated for each approach as follows. RBM-MHC, through the HLA-I classifier, outputs for each peptide a probability to belong to each HLA-I class. MixMHCp2.1 [10–12] models probabilistically data by a mixture of independent models, thus it describes each sequence by a vector of “responsibilities”, describing the probabilities to belong to each cluster (*i.e*., HLA-I class). NetMHCpan4.1 [9] predicts peptide binding values from either the training on eluted data (EL option) or the binding affinity data (BA option) and it estimates from these values presentation scores and percentile rank scores. Low values of percentile rank define binders (following the authors’ recommendations, peptides with percentile rank <2% and <0.5% are considered HLA-I weak and strong binders respectively). Having built the samples from MS data, we considered NetMHC predictions from the EL option (NetMHCpan4.1-EL). We compare the *N^c^* = 6 HLA-I classes by pairs. We consider the probability of belonging to a certain class that each method would assign to “positives” of that class (peptides binding to the respective HLA-I allele) and “negatives” (peptides binding to other alleles): when the classification performance is good, the former has values close to one, the latter has values close to zero. Varying the threshold between false positive and false negative distributions, we build the ROC curve. We take the area under the ROC curve (AUC) as measure of the ability to discriminate the two classes (AUC = 1 means perfect discrimination, AUC = 0.5 means chance). To obtain one cumulative indicator, we average over the AUCs of these pairwise comparisons. For the approaches partially or fully supervised (RBM-MHC and classifier-only), the AUC is measured only from data *not* in the 10% used for the supervised learning step. To further assess classification performance, we looked also at the HLA type of each peptide predicted based on: the highest responsibility value among the *N^c^* values for MixMHCp2.1; the lowest percentile rank for NetMHCpan4.1-EL; the highest class probability estimated by the HLA-I classifier for RBM-MHC. In this way, when comparing peptides of different classes two by two, we can count true positives of classification (*tp*), false positives (*fp*), true negatives (*tn*) and false negatives (*fn*). Once these quantities are defined, additional classification performance indicators are: Accuracy=(*tp + tn*)/(*tp + fp + tn + fn*), Precision=*tp*/(*tp + fp*), Specificity=*tn*/(*tn + fp*), Sensitivity=*tp*/(*tp + fn*). MixMHCp2.1 was run using default options, which include an additional “trash cluster”. When measuring these classification performance indicators, the assignment to the trash cluster is considered among the false negatives. The comparison of classification performance as measured by these indicators is shown in Supplementary Figs. 7B-I.

## Supplementary Information

### Supplementary Figures

**Supplementary Figure 1 (related to Figs. 1B, 2A and STAR Methods).**
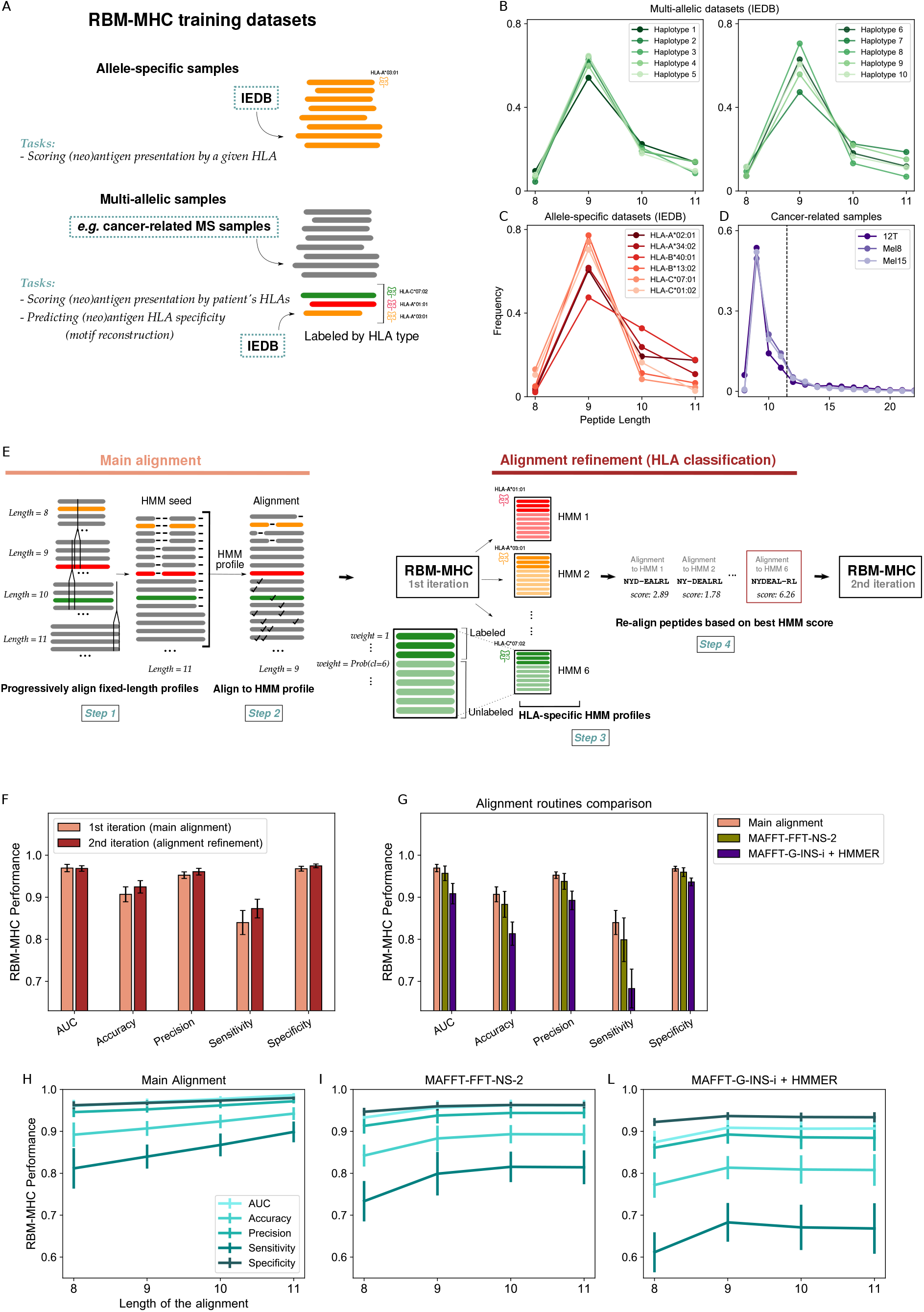
Sequence alignment routine. A: Schematic representation of the types of peptidomic datasets used to train RBM-MHC and the corresponding tasks that RBM-MHC can fulfill. B-D: Sequence length distribution for the different datasets considered in this work: multi-allelic datasets collected from IEDB (B), see Fig. 2C, Supplementary Fig. 7 and Supplementary Table 2; a subset of allele-specific datasets collected from IEDB (E), see Fig. 1D and Supplementary Fig. 5; patient-derived MS samples (D), see Fig. 2D-E and Supplementary Fig. 8. The dashed line in (D) indicates the range of peptide lengths (8-11) considered. E: Schematic representation of the sequence alignment routine, see STAR Methods. In the construction of the main alignment (Step 1 and Step 2) symbol “-” stands for a gap insertion while symbol “ⅴ” stands for the deletion of an amino acid. F: RBM-MHC classification performance indicators evaluated after the 1st and the 2nd RBM-MHC training iterations, as depicted in (E). G: RBM-MHC classification performance (1st iteration only) when the alignment is either performed by our routine, as in (E), or based on MAFFT (MAFFT-FFT-NS-2 and MAFFT-G-INS-i + HMMER, see STAR Methods). In MAFFT-G-INS-i + HMMER, we randomly chose 25% of the sample (hence including sequences of all lengths) as HMM seed. H-L: RBM-MHC classification performance varying the length of the HMM profile in our alignment routine (H) or the length of the alignment by MAFFT-FFT-NS-2 (I) and MAFFT-G-INS-i + HMMER (L) in the range 8-11. (F-L) show the average performance and its standard deviation over the 10 synthetic single-individual samples of Supplementary Table 2.

**Supplementary Figure 2 (related to Figs. 1B, 2A and STAR Methods).**
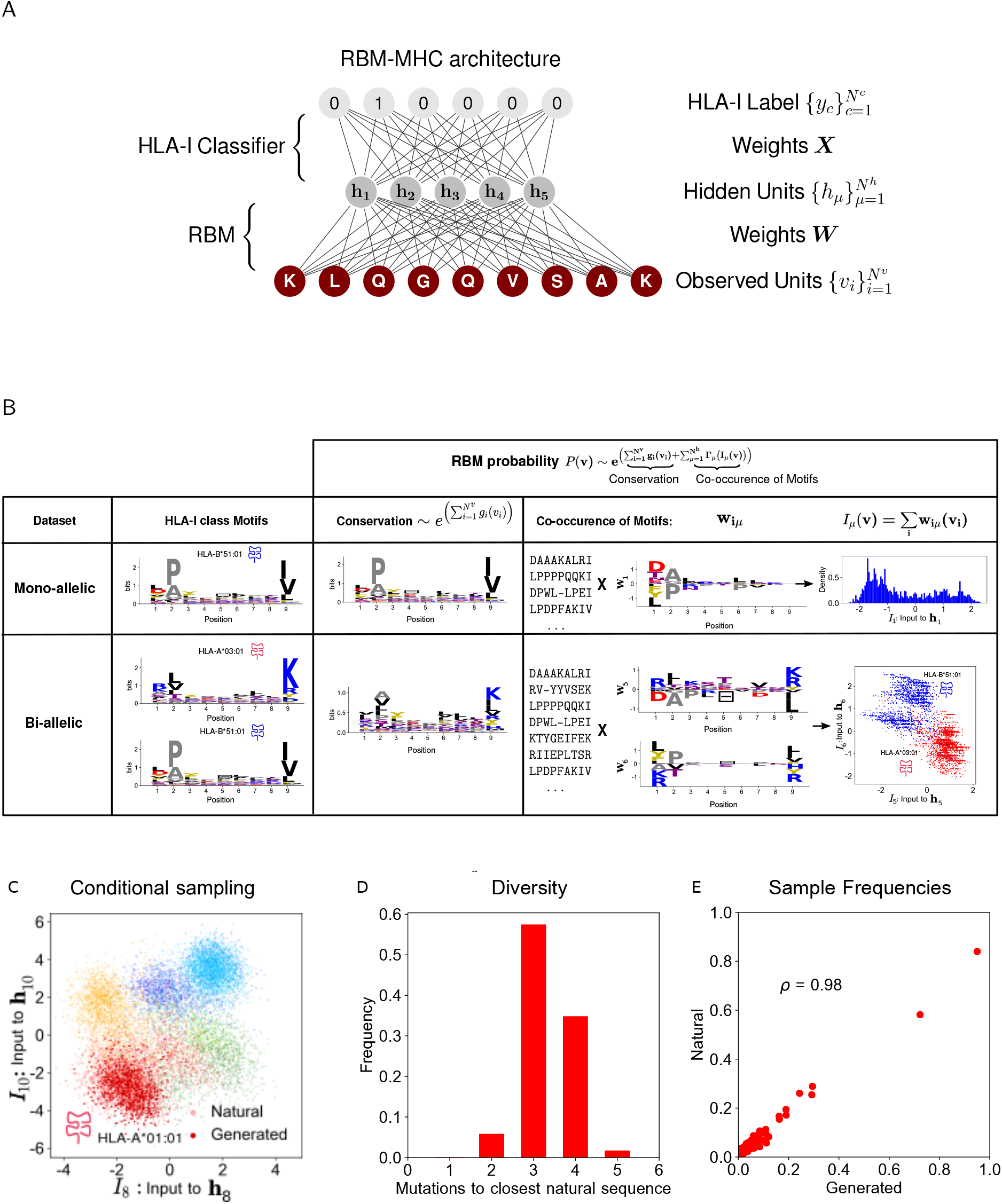
Model’s structure and features. **A: RBM-MHC graphical architecture.** The RBM architecture consists of the layer of observed units ***v*** - the antigen sequences to model - coupled to the layer of hidden units ***h*** by weights ***W***. The HLA-I classifier has a fully connected network of weights ***X*** connecting the layer of inputs to hidden units ***I***(***v***) (RBM data representations) to the output layer of HLA-I classes ***y**_c_* (light gray nodes) formulated in one-hot representation (*e.g*. HLA-I class 2 is encoded by a vector of zeros with the symbol 1 only at position 2). **B: RBM representation of HLA-I specificity: sequence motifs are captured through RBM local potentials (biases) and weights.** The probability of presentation *P*(***v***) inferred by the RBM consists of potentials *g_i_*(*v_i_*), describing single-site amino acid conservation, and terms describing the co-occurrence of motifs through the weights *w_i_μ__*(***v***). In the first row, we consider a IEDB, MS-based dataset of antigens specific to one allele, HLA-B*51:01 (mono-allelic dataset). A RBM model is trained on these data (chosen with 5 hidden units and 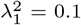, to enhance weights’ interpretability, see Supplementary Fig. 3G). With data of the same HLA-I type, the binding motif is essentially captured by conservation alone (logo representation of *e*^∑_*i*_(*g_i_*(*v_i_*)^, central column). Weights and hidden units embed additional data features (right column). Hidden unit 1 ***h***_1_ encodes the multiple binding modes of HLA-B*51:01 antigens discussed in [12] (see also Fig. 2B): the sequence property “hydrophilic/charge” has a positive projection on ***W***_1_ (and hence positive inputs to ***h***_1_) while the pattern “hydrophobic” has a negative one. In the second row, we train a RBM (10 hidden units and 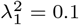) on a bi-allelic dataset, *i.e*. where we added to the mono-allelic dataset IEDB, MS-derived antigen sequences specific to the allele HLA-A*03:01 (a minimal example of a sample mixing antigens of different specificities). In this setting: *(i)* overall conservation does not allow *discrimination* of HLA-I motifs (central column), for which the RBM data representation through weights and inputs to hidden units is necessary; *(ii)* this representation is *robust* with respect to the mono-allelic case. To illustrate this, we have selected sets ***W***_5_ and ***W***_6_ (right column). They distinguish HLA-B*51:01 antigens with sequence patterns underlying alternative binding modes (pattern “hydrophilic/charge” has significant, negative projection on ***W***_5_ and is kept separate from pattern “hydrophobic”, with a significant, positive projection on the other set of weights, ***W***_6_). HLA-B*51:01 sequence patterns are represented as mutually exclusive with respect to the sequence motif characteristic of HLA-A*03:01 (weights of opposite sign). As a result, antigens of different HLA-I types are clearly partitioned into 2 clusters in the space of inputs to ***h***_5_ and ***h***_6_. The 2 subclusters in the HLA-B*51:01 one (blue points) correspond to its alternative binding modes. As conserved sites at position 9 play a role in representing HLA-B*51:01 *vs* HLA-A*03:01 specificity but not in discriminating HLA-B*51:01 binding modes, only in this bi-allelic case weights at that position have a non-negligible value. **Generation of HLA-specific artificial peptides by RBM-MHC**. C: 2-dimensional visualization of the RBM representation space of inputs to hidden units ***I***, as in Fig. 2A. Inputs for antigen sequences with HLA-A*01:01 restriction (light red points) mark a region encoding for this binding specificity. We explore systematically this region by making stochastic moves (points in dark, red color) and by accepting them only if the classifier would predict with high confidence that the sampled antigens specifically bind to HLA-A*01:01 (STAR Methods). In this example, every 100 steps we condition on the current values of inputs and sample 1000 configurations for a total of *M_g_* = 11000 generated sequences. D: Diversity of the generated sequences with the selected HLA-A*01:01 specificity, quantified as number of mutations away from the closest natural sequence of the same specificity. E: Scatter plot comparing positional amino acid frequency of natural and generated antigen sequences with same binding specificity, HLA-A*01:01.

**Supplementary Figure 3 (related to Figs. 1B, 2A and STAR Methods).**
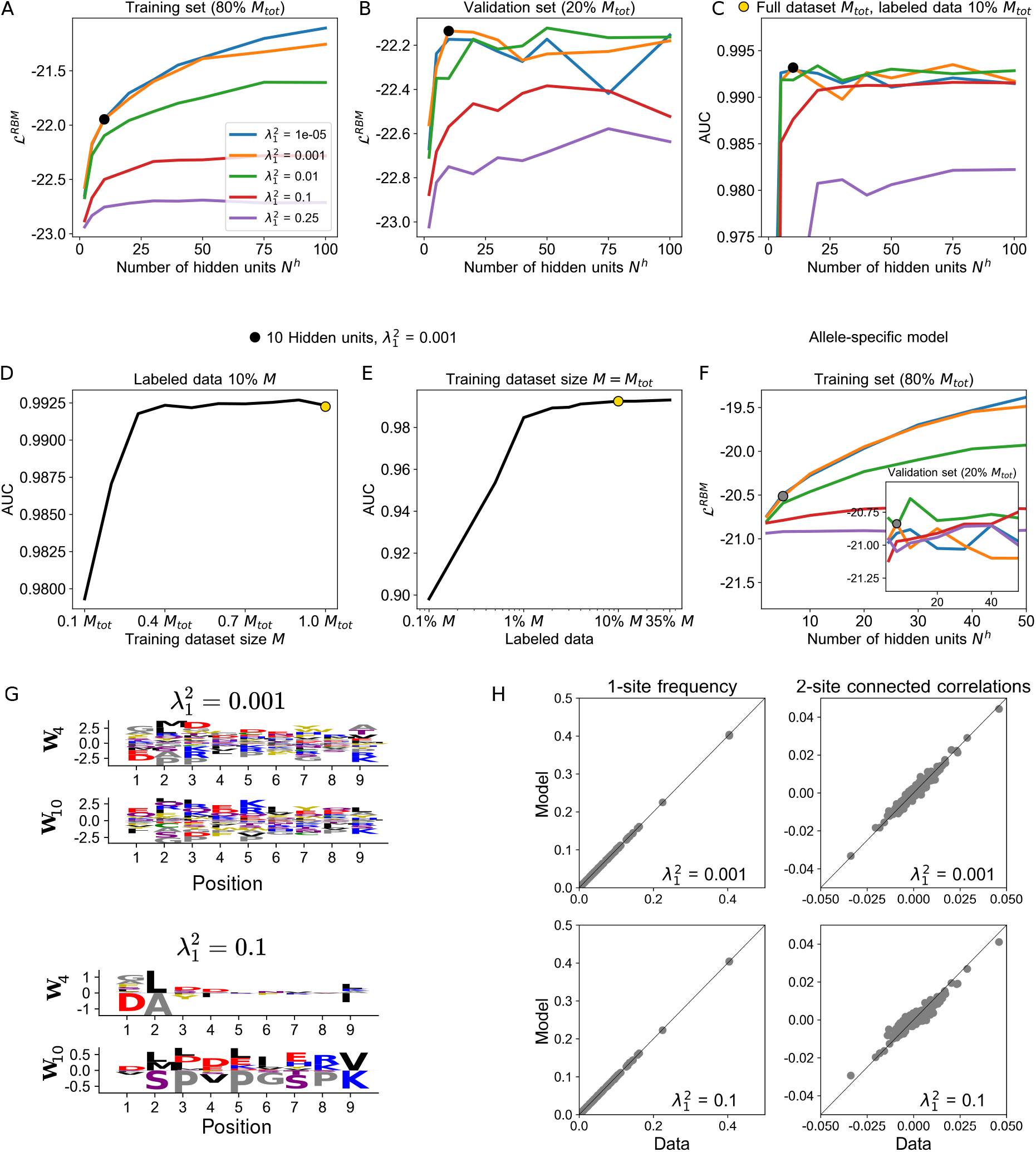
RBM-MHC model selection. RBM average log-likelihood on the training set (A), on the validation set (B) and AUC of HLA-I classification (C) as functions of the number of hidden units *N^h^*, for different values of the regularization strength 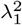. The black dot indicates an optimal combination of parameters chosen for the results in a multi-allelic setting of Figs. 2C-E (*N^h^* = 10, 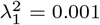). All tests are run on a “synthetic-individual” dataset (Haplotype 10 in Supplementary Table 2, 9-mers only). D: Average AUC of classification as a function of the fraction of data considered as training dataset of the RBM (here *M_tot_* = 9546). E: Average AUC of classification as a function of the RBM training dataset percentage used for the supervised learning of the HLA-I classifier. The light orange dot corresponds to the value chosen for all RBM-MHC results in Figs. 2C-E (*i.e*., *M = M_tot_*, labeled data = 10%M). Curves in (D-E) represent the average over 10 realizations of the training. F: RBM average log-likelihood on the training set and on the validation set (inset) for an allele-specific model (HLA-A*-02:01 model of Fig. 1C,E). The gray dot indicates the parameters chosen for allele-specific models of Figs. 1C-G (*N^h^* = 5, 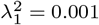). G: Effect of regularization on RBM sequence representation. The RBM learns a set of weights for each hidden unit; in each set, weights have the length of the sequence (9) and for each position there is a weight value for each amino acid appearing at that position that can be either positive or negative (the height of the amino acid letter here gives its magnitude, as in the representation by [16]). Color stands for amino acid chemical/physical properties: red = negative charge (E, D), blue = positive charge (H, K, R), purple = non-charged polar (hydrophilic) (N, T, S, Q), yellow = aromatic (F, W, Y), black = aliphatic hydrophobic (I, L, M, V), green = cysteine (C), grey = other, tiny (A, G, P). Here, we show a comparison of RBM weights (2 sets selected for illustration) for 2 choices of regularization strength 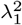 analyzed in (A-E) (training with 10 hidden units, on the “synthetic-individual” dataset corresponding to Haplotype 10 in Supplementary Table 2). As found in [16], at low regularization 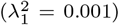 the data representation is more intricate while a higher regularization 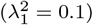, by enforcing sparsity, gives non-zero weights to the amino acids and positions learnt as the most discriminative among “features” (here, HLA-I classes). The first 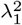 choice however retains more information about data, hence reaches a slightly higher classification performance when combined to the HLA-I classifier (see C); the second 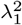 choice results in a more interpretable representation. H: Scatter plots showing that single-site frequency and pairwise connected correlations of the training data are reproduced with high accuracy by the RBM, especially at low regularization (same RBM models as in G).

**Supplementary Figure 4 (related to Fig. 1F and STAR Methods).**
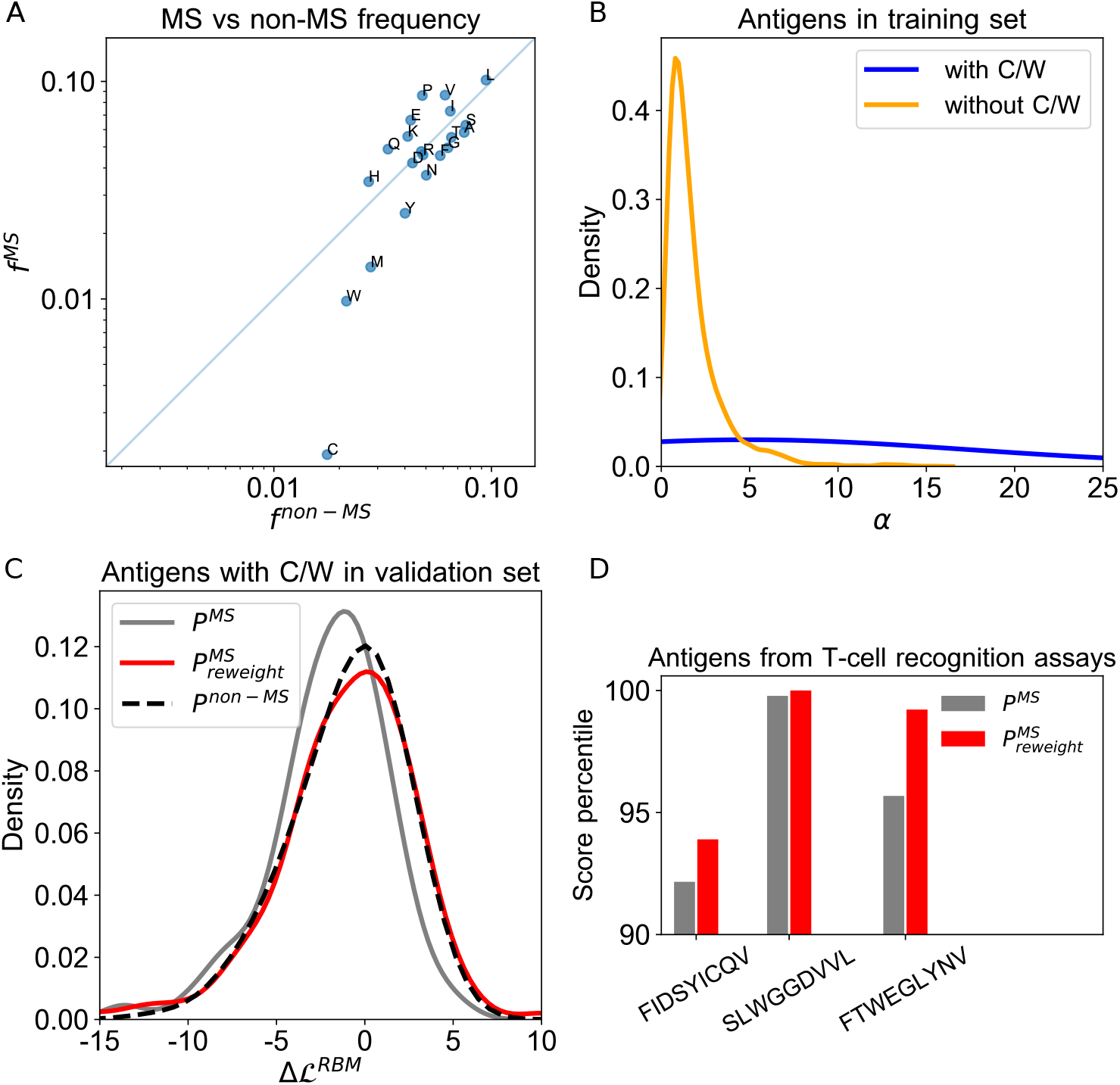
Sequence re-weighting scheme. A: Amino acid frequencies in IEDB antigens detected by MS (*f^MS^*) *vs* all other techniques (*f^non−MS^*): cysteine (C) and tryptophan (W) frequencies are the lowest both for MS and non–MS antigens and in particular their MS estimate has the largest deviation towards very low values compared to the non–MS one, with *f^MS^* ≤ 0.01. B-C: Test of the re-weighting procedure in a RBM model for presentation by HLA-A*01:01 trained on a set of MS data in IEDB (with ligand length in the range of 8-11 residues). B: Distribution of weights αs for ligands in the training dataset with and without C/W. C: Distribution of differences of presentation log-likelihoods defined as 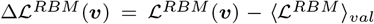, where 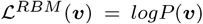, 〈·〉_*val*_ stands for the average over the full validation dataset and ***v*** are sequences in the latter containing C/W. The 3 curves correspond to choosing either *P = P^MS^* (from the RBM model trained on MS data without re-weighting), or 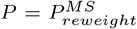 (from the RBM model trained on MS data with re-weighting) or *P = P^non−MS^* (from the RBM model trained on data obtained by techniques different from MS). D: Effect of re-weighting applied to the RBM model for presentation by HLA-A*02:01 of Figs. 1C,E (trained on IEDB, mono-allelic MS-detected ligands of length 8-11 residues). Experimentally validated immunogenic peptides from T-cell recognition assays containing C/W (FIDSYICQV from [45], SLWGGDVVL and FTWEGLYNV from [46]) are assigned an higher RBM presentation score by the re-weighted model 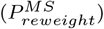 compared to the one without re-weighting (*P^MS^*). Thanks to re-weighting, score percentiles, relative to the score distribution of the 5000 generic peptides considered in Figs. 1C,E, are all improved.

**Supplementary Figure 5 (related to Fig. 1D).**
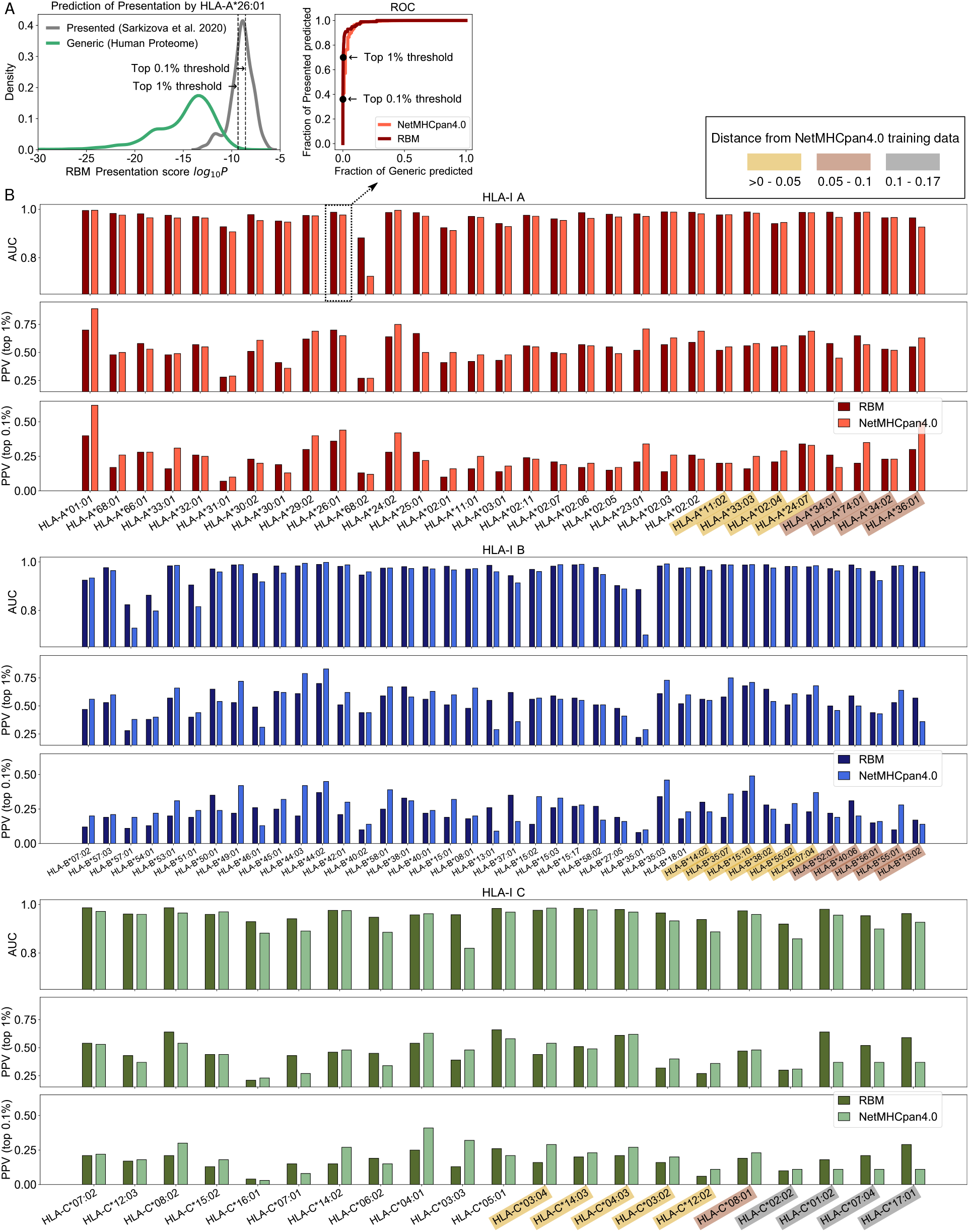
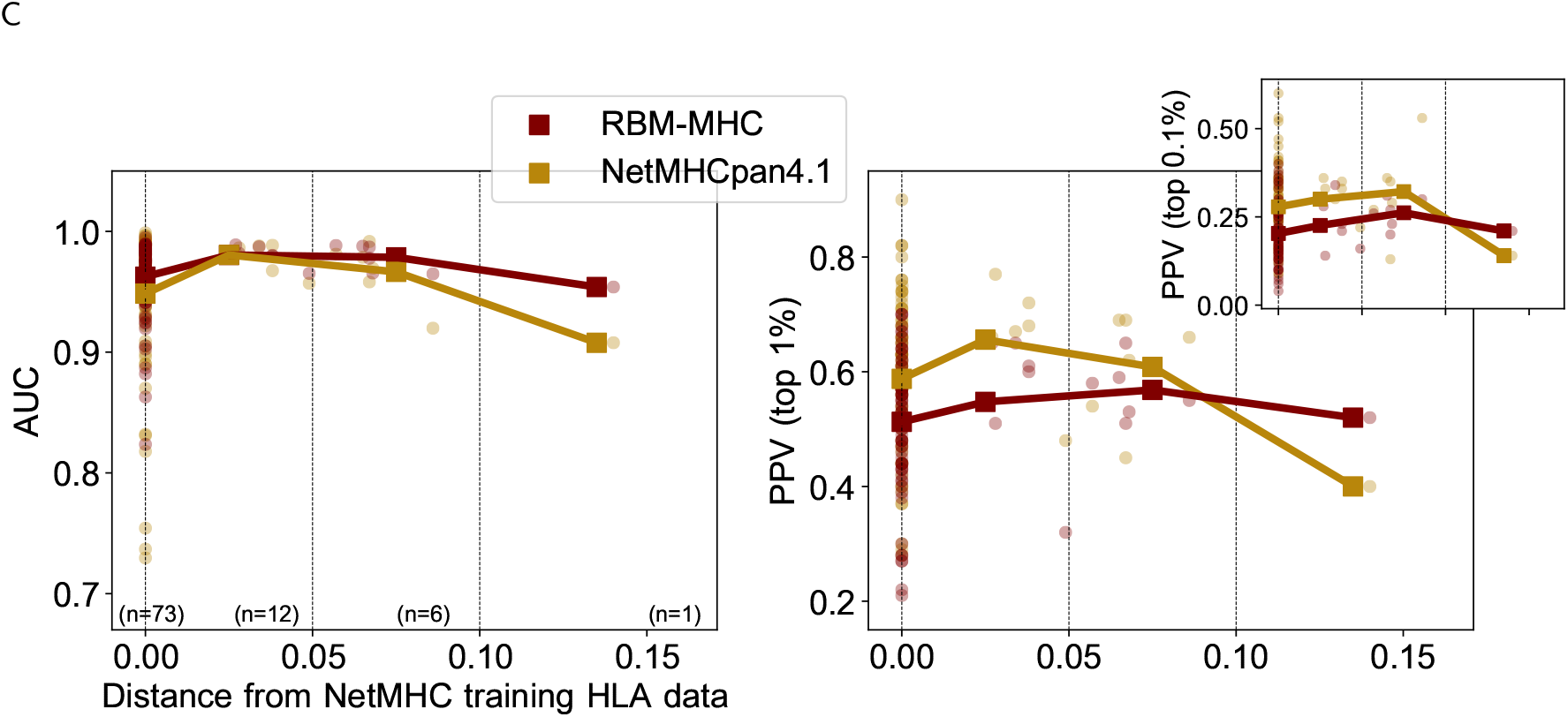
Benchmark of RBM presentation models on mono-allelic datasets from [6]. Comparisons of RBM and NetMHCpan4.0 (with option EL = Eluted) performance at discriminating presented antigens from *n*-fold excess of generic peptides as measured by AUC, PPV at the top scoring 1% of peptides (for *n*=99) and 0.1% (for *n*=999). Subfigure A shows the location of these thresholds in the distribution of RBM presentation scores for an example allele (HLA-A*26:01) along with the full ROC curves given by RBM and NetMHCpan4.0. In B, AUCs and PPVs are shown for each of the 92 HLA-I alleles considered and grouped into 3 subpanels by HLA locus (A,B,C). Colored boxes mark all the alleles for which no HLA data were available in NetMHCpan4.0 training dataset (i.e., the distance to the closest allele in the training set used for prediction is > 0) and the box color denotes the interval of distance values. C: Same plot as in Fig. 1D showing the comparison of RBM to NetMHCpan4.1. NetMHCpan4.1 performance is improved with respect to RBM in terms of PPVs, except for the rarest allele, but AUCs are still completely comparable.

**Supplementary Figure 6 (related to Fig. 1F-G).**
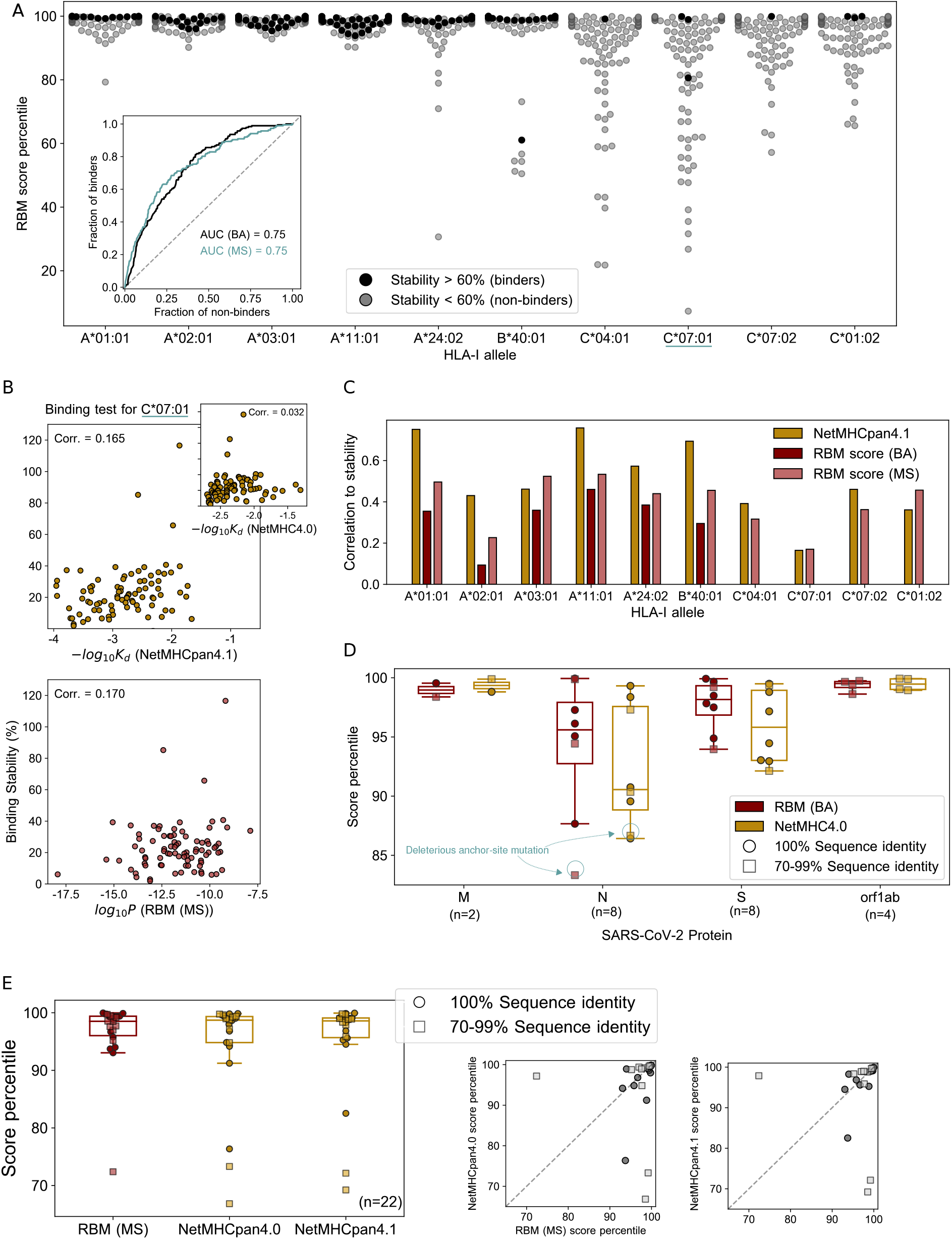
**A-C: RBM prediction of SARS-CoV-2 epitopes tested *in vitro*.** A: RBM score percentile of putative SARS-CoV-2 ligands predicted by NetMHC (NetMHC4.0/NetMHCpan4.0) for 10 HLA-I alleles (94 ligands per allele), divided into “binders” and “non-binders” on the basis of the binding stability assigned *in vitro* by [24] (respectively > 60% and < 60% of reference values). RBM models were trained on binding assay (BA) data for HLA-A and HLA-B, on mass spectrometry (MS) data for HLA-C (see Fig. 1F and STAR Methods). The average RBM score percentile for binders is consistently above 98 for each HLA-I allele. We quantify the RBM ability to distinguish binders from non-binders by the Receiver Operating Characteristic curve (inset), which describes, as a function of a score percentile threshold, how the fraction of binders and non-binders is recovered by the RBM score. Its Area Under the Curve (AUC) is significantly above the random expectation of 0.5. As a comparison, we added the AUC obtained using RBM models all trained on MS data, giving a comparable performance. An unbiased comparison to NetMHC is here difficult because peptides were pre-selected ahead of the experiments from the top 1% ranks estimated by NetMHC4.0/NetMHCpan4.0. B: We consider the peptides tested for an example of HLA-C allele in (A), C*07:01. The scatter plots show the trend of binding stability measured in Ref. [24] *vs*: the RBM score *log*_10_*P* (bottom), using the same RBM model as in (A); the NetMHCpan4.1/NetMHC4.0 score (top), given by −*log*_10_*K_d_*, where *K_d_* is the predicted binding affinity in [nM]. NetMHCpan4.1 was run with the option BA = Binding Affinity. The Pearson correlation (corr.) between the quantities on the *x*- and *y*-axis is explicitly reported. C: Same plot as Fig. 1F where NetMHC scores are computed by the NetMHCpan4.1 version and the correlation is estimated as illustrated in (B). **D-E: RBM scores for SARS-CoV-2 homologs of tested, dominant SARS-CoV epitopes.** D: Same data as in Fig. 1G, where epitopes are divided by SARS-CoV-2 protein, with S = Spike glycoprotein, E = Envelope protein, M = Membrane glycoprotein, N = Nucleocapsid phosphoprotein. The lowest RBM score percentile (consistent with NetMHC4.0 one, see also in Fig. 1G) corresponds to an anchor-site mutation of the SARS-CoV homolog expected to be deleterious, as it leads from a highly abundant amino acid at position 9 to a low-frequency one. E: Same figure as in Fig. 1G with scores obtained by MS-trained RBM models. Here the comparison is made with NetMHC algorithms trained on MS datasets, *i.e*. NetMHCpan4.0 and NetMHCpan4.1 (run with the option EL = Eluted).

**Supplementary Figure 7 (related to Fig. 2C).**
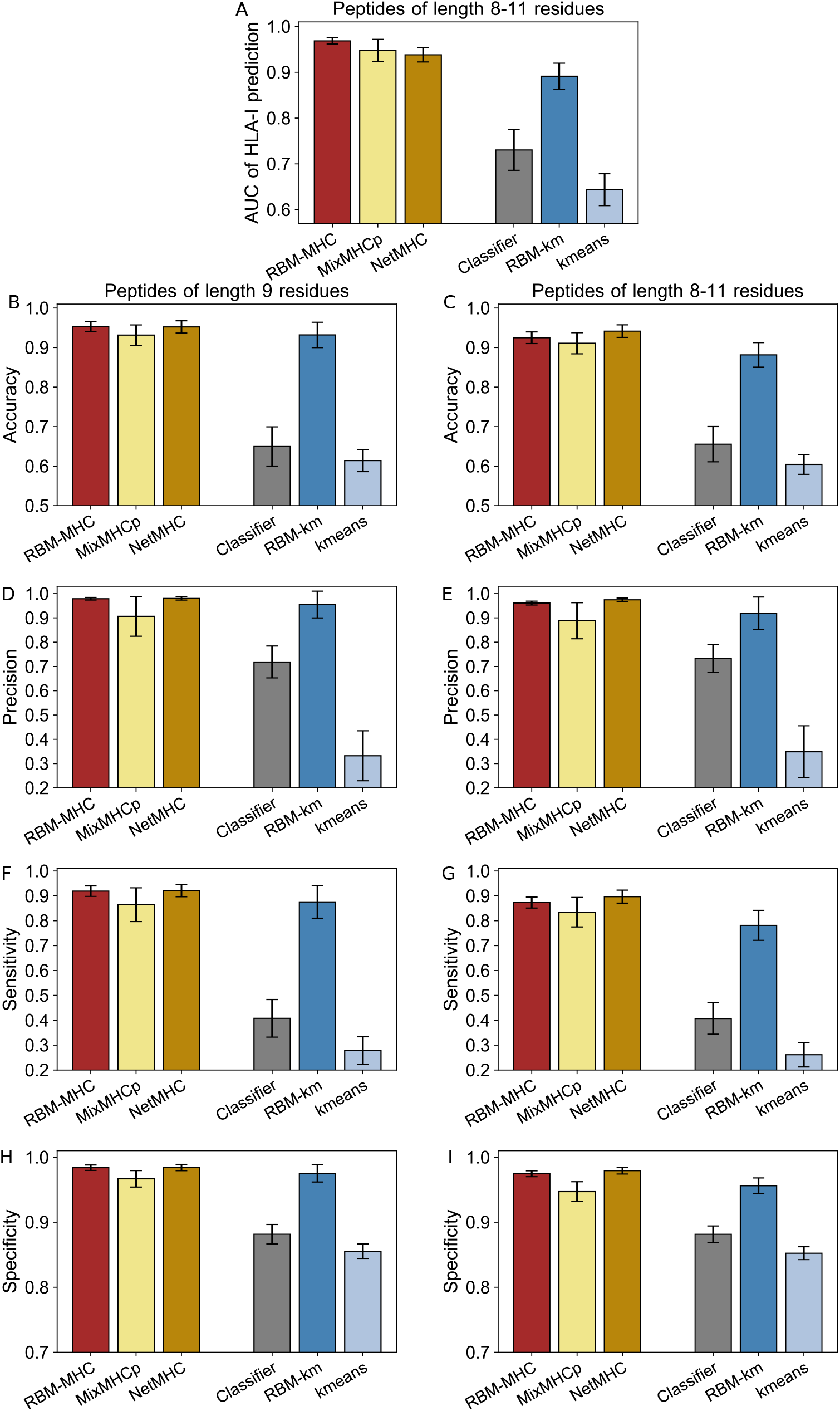
Performance at predicting antigen HLA association in “synthetic-individual” samples. A: Same plot as in Fig. 2C for datasets containing peptides of length 8-11 residues, where a sequence alignment step is required (see STAR Methods). With the same representation as in Fig. 2C and (A), we show: average accuracy (B,C), precision (D,E), sensitivity (F,G), specificity (H,I) (see STAR Methods). Classification performance indicators are measured separately in datasets containing only peptides of length 9 (B,D,F,H) and in datasets containing peptides of length 8-11 (C,E,G,I).

**Supplementary Figure 8 (related to Fig. 2D-E).**
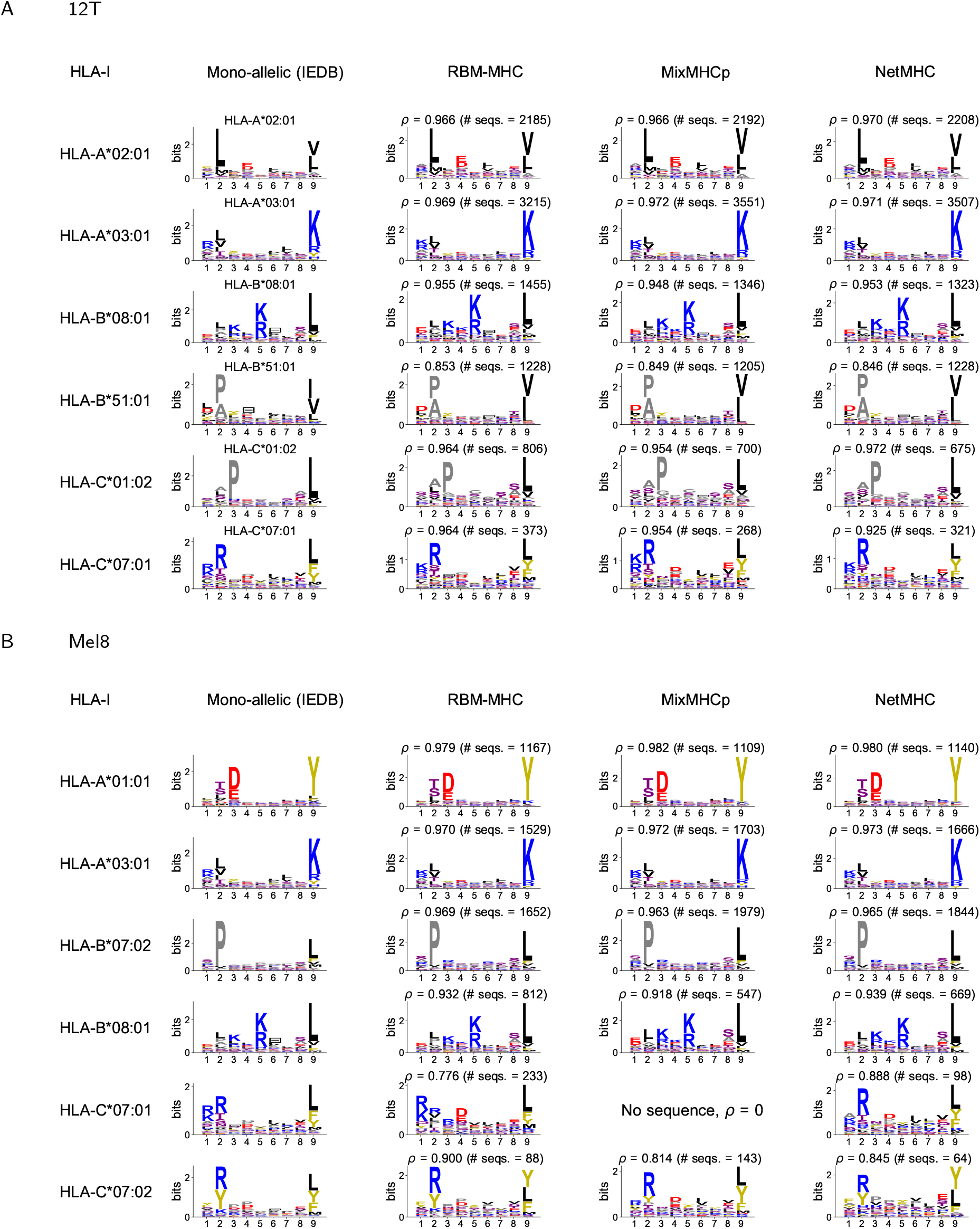
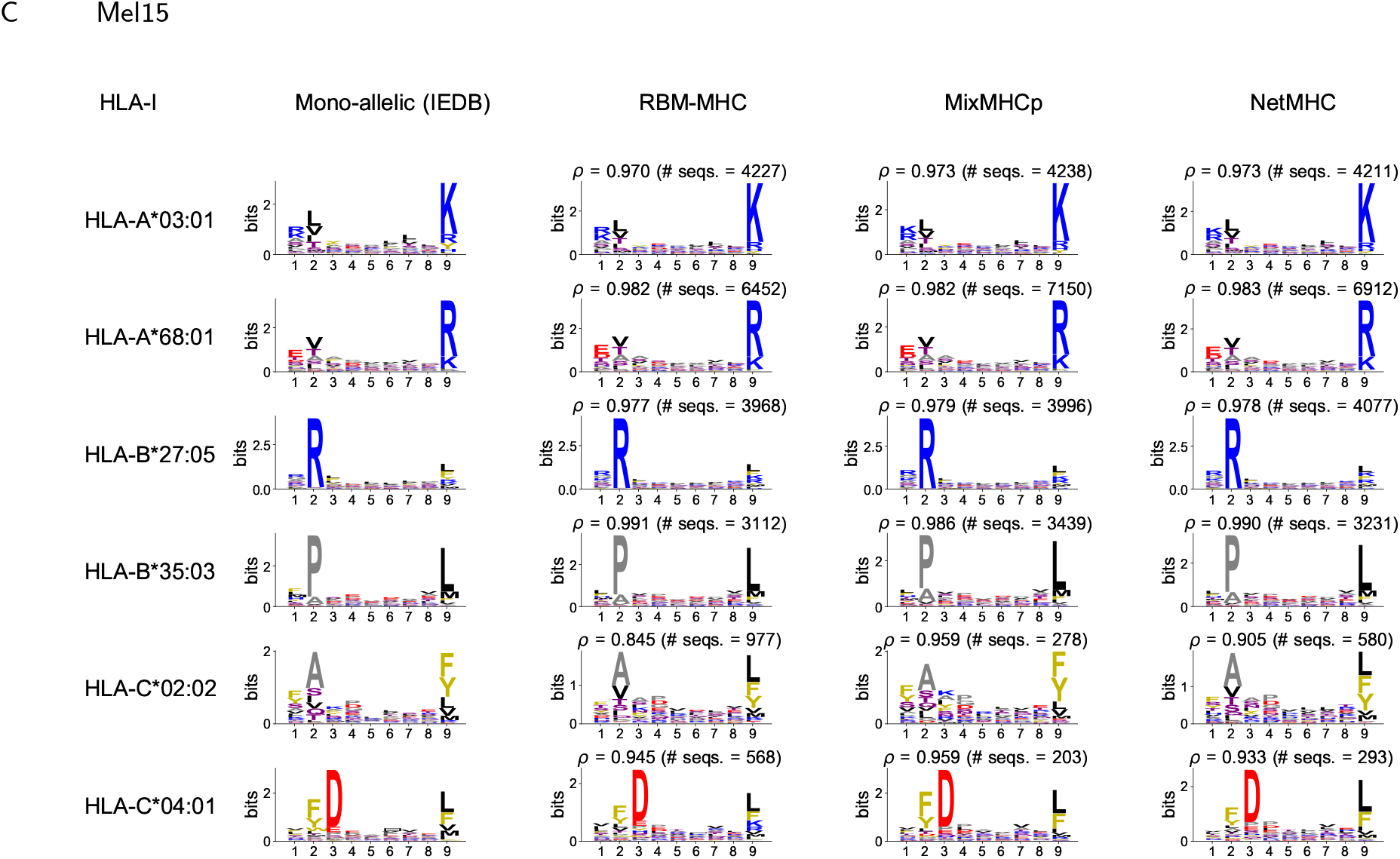
Sequence motif reconstruction in multi-allelic samples. A: Sample 12T from [26, 27], sample size = 9262 sequences (restricted to length 8-11); B: sample Mel8 from [31], sample size = 5481 sequences (restricted to length 8-11); C: sample Mel15 from [31], sample size = 19304 sequences (restricted to length 8-11). Sequence motifs recovered by RBM-MHC are compared to the ones from mono-allelic data in IEDB and the ones recovered by MixMHCp2.1 and NetMHCpan4.1-EL in the same sample. The quality of motif reconstruction is measured by *ρ*, the coefficient of correlation between single-site amino acid frequencies of the mono-allelic IEDB motif and of the predicted one. The average *ρ* over the 6 HLA-I motifs recovered by the 3 methods is compared in Fig. 2E. Motifs of length 9 in mono-allelic data are obtained by aligning sequences in each HLA-I class separately, using standard alignment routines as described in STAR Methods. When comparing these motifs to the ones predicted by the different methods, we keep the alignment step consistent with the choice made for mono-allelic data, *i.e*. for each method we align separately, applying the same standard alignment routines, the sequences assigned to the same HLA-I class.

### Supplementary Tables

**Supplementary Table 1 (related to Fig. 1F).**
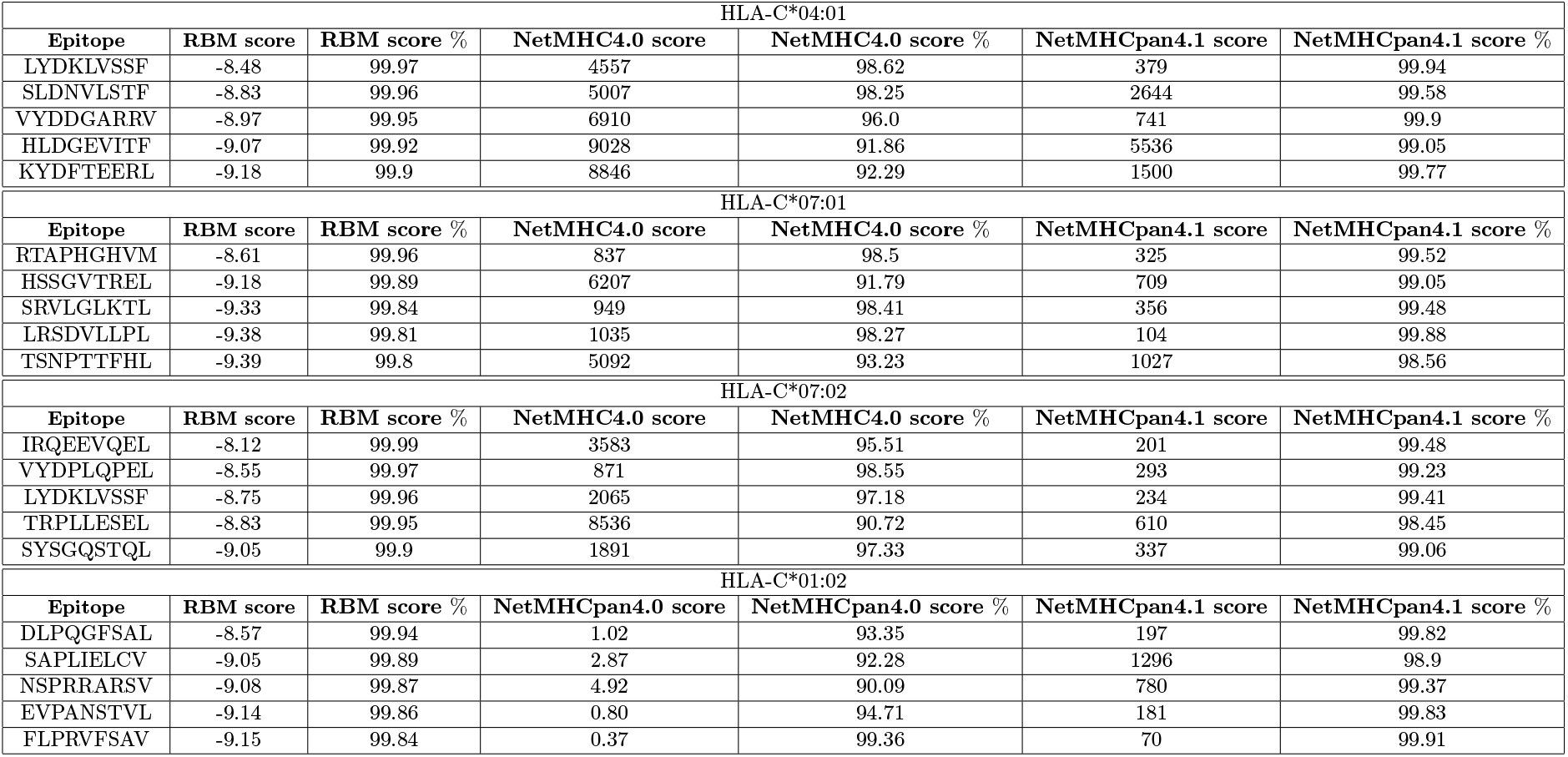
Cross-validation of SARS-CoV-2 epitopes by RBM and NetMHCpan4.1. Lists of 5 SARS-CoV-2 peptides for each HLA-C considered in [24] along with their scores from RBM, NetMHC4.0/NetMHCpan4.0 (the methods used in [24] to rank epitopes ahead of the experimental test), NetMHC-pan4.1 and the respective score percentiles (relative to the full set of SARS-CoV-2 9-mers). The score from RBM gives the peptide log-likelihood (log base 10), scores from NetMHC4.0 and NetMHCpan4.1 (which we run with BA option) represent the peptide binding affinity (expressed in [nM]) while the score from NetMHCpan4.0 (which was run with EL option in Ref. [24]) is a rank percentile. The 5 peptides per allele were chosen as examples of candidate binders predicted by RBM that were not selected among the top-scoring ones by NetMHC4.0/NetMHCpan4.0 and hence whose stability was not tested in Ref. [24]. RBM predictions for these peptides are corroborated by NetMHCpan4.1, which inserts all these peptides among its predicted “strong binders”. The agreement between RBM and NetMHCpan4.1 is clear from the high score percentiles assigned by both methods (columns 3 and 7).

**Supplementary Table 2 (related to Fig. 2C and Supplementary Fig. 7, provided as a separate Excel file). Detailed benchmark of RBM-MHC prediction performance**. Allelic composition of “synthetic-individual” samples retrieved from IEDB (see STAR Methods) and measures of AUC, accuracy, precision, sensitivity, specificity for each sample.

**Supplementary Table 3 (related to Fig. 2D-E).**
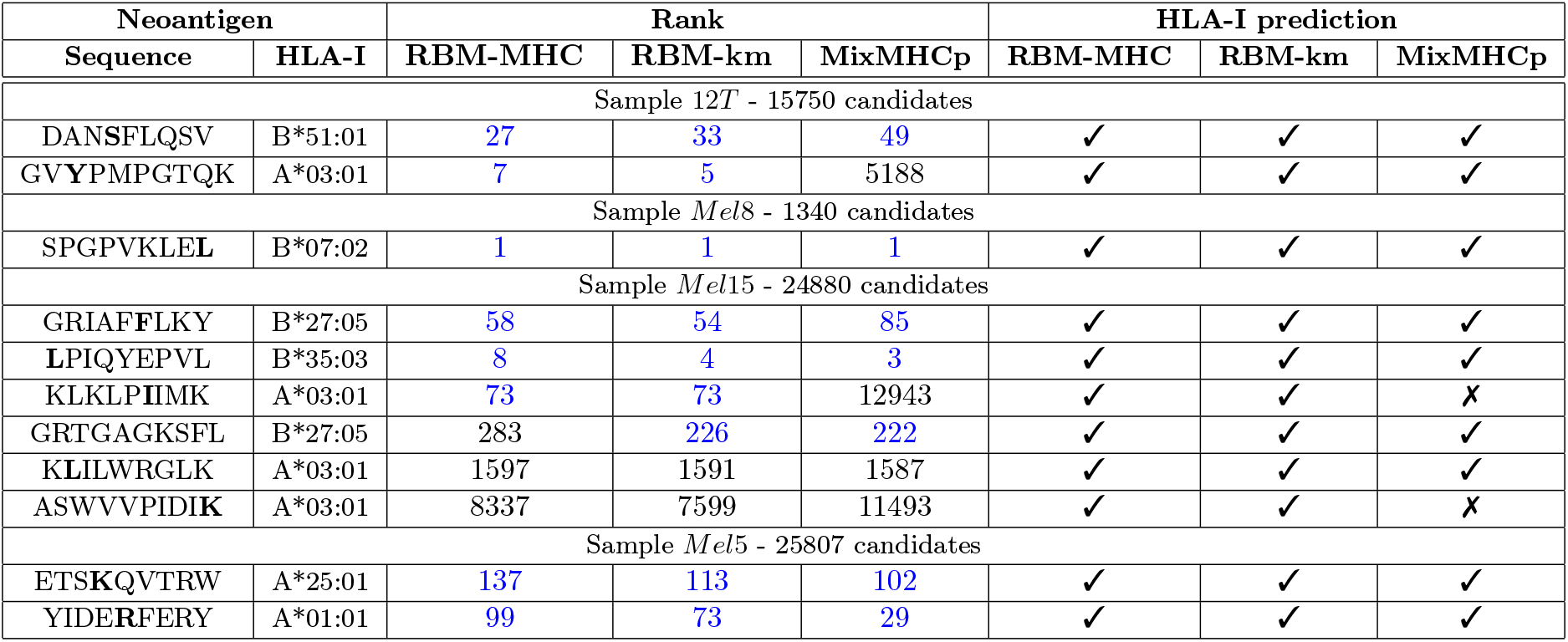
Ranking and HLA-I restriction prediction for neoantigens from cancer-related samples. (12T from [26, 27] and Mel8, Mel5 and Mel15 from [31]). The mutated residue in each neoantigen sequence is marked as a bold letter. All neoantigen candidates (whose number is reported in the header) were ranked according to the presentation scores assigned by RBM-MHC (see Eq. 9 in STAR Methods) in decreasing order. RBM-MHC predictions are compared to the same ranking by the two unsupervised approaches, MixMHCp2.1 and RBM-km (both trained on the same single-patient dataset). For each method, we have highlighted in blue ranks in the top 1%, *i.e*. higher than the 99% of the full set of neoantigen candidates. This comparison shows that: the RBM-km attains a performance comparable to RBM-MHC (8/11 ranked among top 1%, expected HLA restriction systematically retrieved), emphasizing the key role of RBM modeling for predictive power; MixMHCp provides accurate predictions (4/11 times gives higher rank than RBM-MHC) but in a less robust way, missing 2 neoantigens among the top 1% and 2 HLA-I restrictions that RBM-MHC and RBM-km assess well. The same ranks estimated by NetMHCpan4.0 and MixMHCpred, available respectively in [8, 11] (apart from the neoantigen GVYPMPGTQK, detected in a more recent analysis [27]), are overall higher than RBM-MHC ones, largely due to training on large-scale, optimized datasets. However, NetMHCpan4.0 fails at placing in the top 1% the same 3 neoantigens for which the RBM-MHC fails, and so does MixMHCpred for 2 of them. Hence, the RBM-MHC bears the comparison to these methods, given that it is trained on a moderate-size, patient-derived dataset instead of a large-scale, optimized dataset as NetMHCpan4.0 and MixMHCpred.

## Notes

### Competing Interest Statement

The authors have declared no competing interest.

